# Exploring the Potential of Structure-Based Deep Learning Approaches for T cell Receptor Design

**DOI:** 10.1101/2024.04.19.590222

**Authors:** Helder V. Ribeiro-Filho, Gabriel E. Jara, João V. S. Guerra, Melyssa Cheung, Nathaniel R. Felbinger, José G. C. Pereira, Brian G. Pierce, Paulo S. Lopes-de-Oliveira

## Abstract

Deep learning methods, trained on the increasing set of available protein 3D structures and sequences, have substantially impacted the protein modeling and design field. These advancements have facilitated the creation of novel proteins, or the optimization of existing ones designed for specific functions, such as binding a target protein. Despite the demonstrated potential of such approaches in designing general protein binders, their application in designing immunotherapeutics remains relatively unexplored. A relevant application is the design of T cell receptors (TCRs). Given the crucial role of T cells in mediating immune responses, redirecting these cells to tumor or infected target cells through the engineering of TCRs has shown promising results in treating diseases, especially cancer. However, the computational design of TCR interactions presents challenges for current physics-based methods, particularly due to the unique natural characteristics of these interfaces, such as low affinity and cross-reactivity. For this reason, in this study, we explored the potential of two structure-based deep learning protein design methods, ProteinMPNN and ESM-IF, in designing fixed-backbone TCRs for binding target antigenic peptides presented by the MHC through different design scenarios. To evaluate TCR designs, we employed a comprehensive set of sequence- and structure-based metrics, highlighting the benefits of these methods in comparison to classical physics-based design methods and identifying deficiencies for improvement.

## 1 Introduction

T cell receptors (TCRs) are heterodimeric receptors located on the T cell surface, specialized in recognizing and binding to antigenic peptides presented by the major histocompatibility complex Class I (MHC-I) or Class II (MHC-II). The peptide recognition by the TCR enables T cells to identify foreign proteins from infectious organisms and viruses, as well as altered proteins in tumor cells, and respond accordingly. The canonical binding of TCR to peptides presented by the MHC (pMHC) canonically involves the TCR complementary-determining region (CDR) loops (CDR1, CDR2, and CDR3) of both heterodimeric chains (α and β chains, for instance). The key specificity in TCR binding primarily residues in the interaction of CDR3 loops, particularly the CDR3β, which is the most variable region of the TCR as a consequence of random rearrangements of variable (V), diversity (D), and joining (J) gene segments [1].

Given the pivotal role of T cells in orchestrating the adaptive immune response, substantial efforts have been employed to genetically engineer T cells capable of expressing TCRs designed to recognize specific antigens. This strategy, known as TCR-engineered T cell (TCR-T) therapy, has emerged as a notable route for immunotherapy, particularly in the context of cancer [2, 3, 4]. When compared to other immune response-based therapies like antibody and chimeric antigen receptor T cell (CAR-T) therapies, TCR-based approaches offer distinct advantages centered on their capacity to efficiently recognize a wide universe of antigens from various subcellular compartments, extending beyond surface-expressed targets [5]. Over the past few years, no fewer than 25 phase I/II clinical trials involving TCR-T cell therapy, targeting diverse epitopes, have been reported for the treatment of solid tumors, emphasizing the significant efforts in this field [4]. In 2022, the FDA approved tebentafusp, the first TCR-based therapy to target the gp100 epitope, used to treat HLA-A*02:01-positive adult patients with unresectable metastatic uveal melanoma [6]. Promising results have also been observed in clinical trials for high-affinity engineered TCRs such as afamitresgene autoleucel, which targets the MAGE-A4 melanoma-associated antigen presented by HLA-A*02 [7].

Despite the advancements in TCR-T therapy, the intricate and still poorly understood mechanism of antigen recognition by TCRs and subsequent T cell activation present challenges to the rational design of TCRs for therapeutic applications. Unlike antibodies, which undergo an affinity maturation process involving somatic hypermutations to enhance affinity for the target, natural TCRs typically exhibit low affinities in the micromolar range [8]. In certain scenarios, such as therapies targeting tumor-associated antigens, the practical application of natural TCRs is limited due to the elimination of T cells recognizing self-antigens during thymic selection [4, 9]. To address this limitation, enhanced affinity TCRs can be generated using various experimental techniques, such as phage display [10, 11, 12, 13]. However, extreme increases in TCR affinity to supraphysiological levels are associated with impaired T cell activation [14]. Furthermore, substantial increases in TCR affinity for a specific target may lead to increased affinity for other unintended targets, resulting in cross-reactivity. For instance, cross-reactivity with self-antigens can result in off-target effects, as observed in a notable case where the use of a TCR with improved affinity for binding the MAGE-A3 antigen from cancer cells induced cardiac toxicity in treated patients due to the high similarity of this peptide to a self-peptide from the cardiac Titin protein [15, 16]. Given the potential drawbacks associated with supraphysiological affinities, the development of TCRs with low micromolar affinities is advocated as a viable solution.

In this context, solved 3D atomic structures or molecular models of TCRs in complex with antigenic peptides bound to the MHC provide valuable details about the TCR recognition mechanism and can serve as the basis for the rational design of TCRs through computational tools. By leveraging these 3D structures, a refined balance between affinity and specificity can be achieved by focusing the TCR design toward peptide contact regions rather than the MHC, thereby mitigating cross-reactivity [17]. Structure-based computational design approaches utilizing physics-based scoring functions have been successfully employed in enhancing the affinity of the DMF5 TCR against melanoma-associated antigens, yielding a remarkable 400-fold increase in binding affinity [18]. However, it is noteworthy that these scoring functions face challenges in generalizing across various TCR:pMHC complexes [19], necessitating calibrations to enhance accuracy despite their significance in TCR optimization.

On the other hand, the growing availability of general protein 3D structures and sequences, alongside recent advancements in machine and deep learning techniques, has significantly accelerated the accurate prediction of protein amino acid sequences capable of folding into a specific backbone structure. Within this context, ProteinMPNN is revolutionizing the protein design field by leveraging the power of Graph Neural Networks and Message-Passing algorithms, alongside an extensive training set of protein structures, demonstrating remarkable capabilities in designing entirely novel proteins [20]. Additionally, the ESM inverse fold (ESM-IF) model, trained on 12 million AlphaFold2-modeled protein structures combined with experimentally determined ones, using Graph Neural Networks with Geometric Vector Perceptron layers, has shown promising results in predicting protein sequences that fold into a predetermined backbone structure [21, 22]. While the applicability of these models has been demonstrated in the design pipeline of protein binder interfaces [20, 21, 23], their potential in TCR design remains unexplored.

In this study, we extensively explored the use of ProteinMPNN and ESM-IF for structure-based TCR interface design, encompassing various design scenarios and comparing them with a widely used physics-based method, the Rosetta Deisgn [24]. To assess the efficacy of these deep learning models, we compared the designs with known TCRs by basing the design process on experimentally solved TCR structures in complex with the pMHC. We employed a comprehensive and orthogonal set of evaluation metrics, beginning with sequence-based metrics that included the rate of native sequence recovery and physicochemical similarity at interface hotspot positions. As a complementary strategy for evaluating the designs, we employed a structure-based metric by modeling the design sequences with TCRModel2 [25] and assessing the confidence of the models. The TCR designs were further evaluated using Rosetta physics-based metrics to assess interface quality and complex stability. Finally, to compare the binding affinity of the designed complexes with native TCRs, we employed robust protocols of Molecular Dynamics (MD) simulations coupled with Molecular Mechanics Poisson-Boltzmann Surface Area (MM/PBSA) free energy calculations. To ensure accurate binding affinity prediction, this protocol underwent benchmarking against a curated dataset of TCR:pMHC structures with experimental binding affinity measurements.

We found that ProteinMPNN and, particularly, ESM-IF demonstrated a remarkable ability to recover native TCR interface amino acid sequences compared to a classical physics-based method, successfully reproducing the amino acid composition of this interface. Importantly, this recovery was not solely based on the backbone or intra-TCR contacts but heavily relied on specific interactions with the pMHC. Alongside high sequence recovery rates, the TCR designs exhibited modeling and energetic scores comparable to native structures. Despite the majority of designs showing similar predicted binding affinities, we identified some designs with improved affinity compared to the native TCR.

We aim for this study to serve as a guiding framework for the structure-based design of TCRs using deep learning and thus provide the basis for the computational engineering of specific TCRs targeting antigenic peptides presented by the MHC, such as tumor and viral antigens.

## 2 Results

### 2.1 Deep learning-based methods outperforms physics-based methods in recovering TCR native sequences

To assess the capability of deep learning and physical-based methods in designing TCR sequences, based on 3D structures, for targeting specific pMHC complexes, we curated a structural dataset consisting of non-redundant TCR:pMHC complexes (32 MHC-I complexes and 6 MHC-II complexes; see Methods section for details). These complexes served as native reference structures for comparison with the generated designs.

As a first TCR design scenario, we focused on designing the TCR CDR3 positions (α and β chains) at the interface with the pMHC (Figure 1A). Considering this scenario, on average, we designed 10 and 11 positions of the CDR3s per each TCR:pMHC-I and MHC-II test case, respectively (Figure S1).

**Figure 1:**
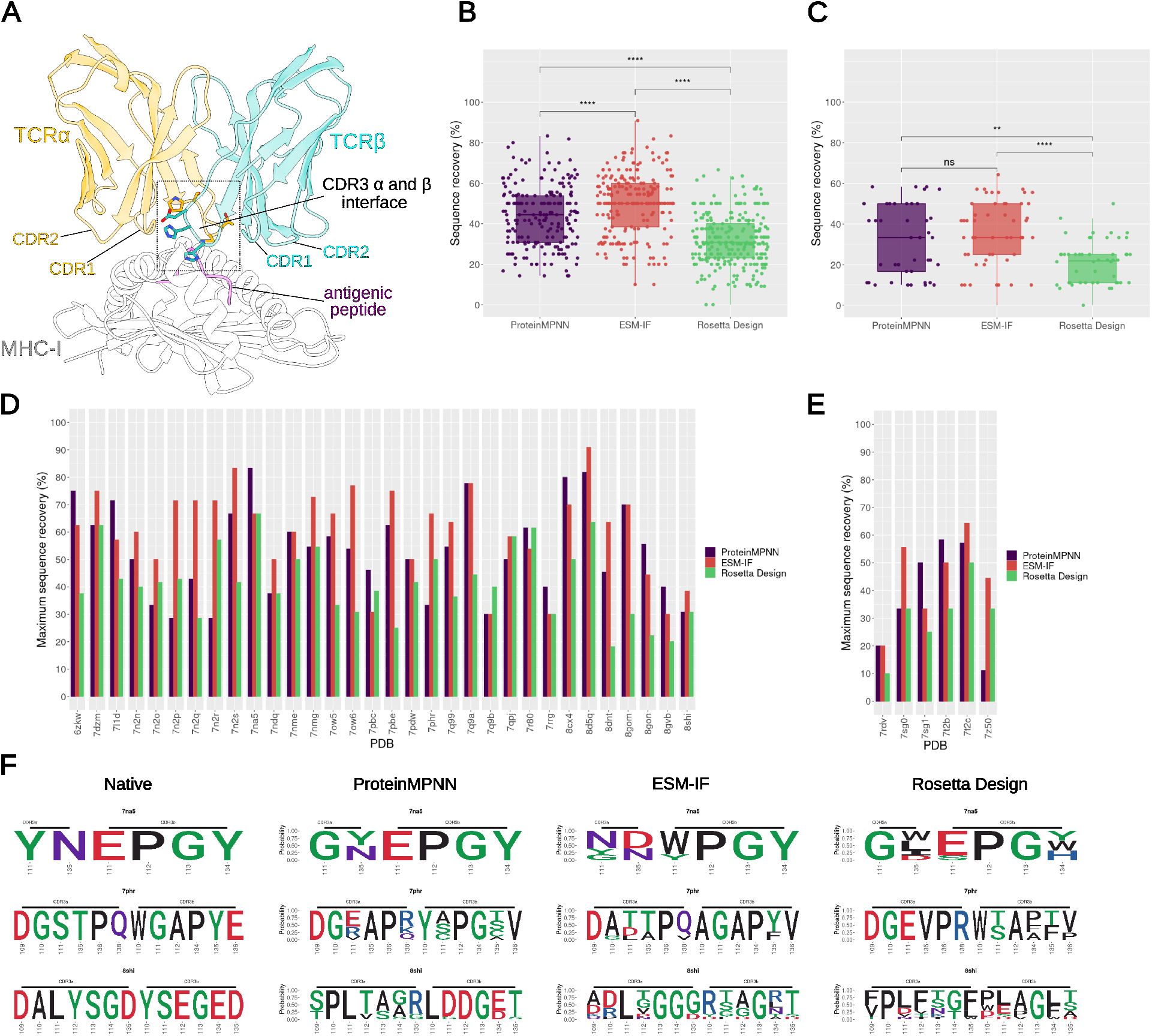
Sequence recovery analysis of interface CDR3s amino acids in designs with ProteinMPNN ESM-IF or Rosetta Design (InterfaceDesign2019 protocol). **(A)** Representative structure of a TCR:pMHC complex (PDB ID: 7nme). The TCR variable and MHC chains were trimmed to just include components spatially related to the interface. The interface is indicated and the CDR3s amino acids composing the interface are shown as sticks. **(B)** Percentage of sequence recovery per method considering all MHC-I test cases. Each point represents a unique design sequence from a test case. For each test case, a total of 10 designs were generated by each method, but redundant designs were removed from the plot. For ProteinMPNN we employed a temperature sampling of 0.1, whereas for ESM-IF a temperature sampling of 0.2 was used (see methods). Statistical two-sample pairwise comparison between methods were performed using Mann-Whitney test with the R ggpubr package. Significance is indicated above each box plot (**** and ** correspond to a p-value below 0.0001 and 0.01, respectively, while ‘ns’ means no significance). **(C)** same as (B), but for MHC-II. **(D)** Maximum sequence recovery obtained for each MHC-I test case and **(E)** for MHC-II test cases. **(F)** Sequence logo of three MHC-I test cases: 7na5, 7qhr, and 8shi. Each row of the panel corresponds to a specific test case and each column corresponds to the design method applied. The first column presents the native amino acids.

Our initial assessment of design success involved comparing the identity of amino acids generated at the designed positions to the native amino acids, computing the percentage of sequence recovery for each test case. A critical parameter that affects sequence design, especially the diversity of generated sequences, in deep learning-based methods is temperature sampling. For ProteinMPNN, we employed a temperature of 0.1, which was used by its developers and others [20, 23]. In our assessments, this temperature achieved the highest sequence recovery while producing higher sequence entropy and uniqueness than lower temperatures (Figure S2). To ensure comparability, we chose a temperature (T = 0.2) for ESM-IF that resulted in sequences with similar sequence entropy to those from ProteinMPNN at T = 0.1. These temperatures for ProteinMPNN and ESM-IF were also consistent in terms of the percentage of uniqueness (Figure S2).

On average, ProteinMPNN and ESM-IF achieved sequence recovery percentages of 43.9% and 50.1%, respectively, for MHC-I, and 32.0% and 35.6% for MHC-II cases, with each test case generating 10 designs (Figure 1B and Figure 1C). Sequence recovery rates per target are presented in Figure S1. Notably, both deep learning-based methods, ProteinMPNN and ESM-IF, surpassed the sequence recovery of designs generated by Rosetta FastDesign protocol [24], with sequence recovery percentages of 31.7% and 19.6% for MHC-I and MHC-II cases, respectively. In 66% and 84% of test cases (including both MHC-I and MHC-II), ProteinMPNN and ESM-IF, respectively, generated designs with superior sequence recovery compared to Rosetta Design. As an alternative to the default Rosetta FastDesign approach, we also explored protocol variations. This included generating 1000 designs and calculating the sequence recovery over the top 10 designs ranked by Rosetta energy terms. However, we observed consistent trends across these variations (Figure S3).

When comparing ProteinMPNN and ESM-IF, the latter generated designs with higher sequence recovery than ProteinMPNN in 55% of cases, while ProteinMPNN generated higher sequence recovery rates in 29% of cases (Figure 1D and 1E). On average, ProteinMPNN, ESM-IF, and Rosetta FastDesign achieved maximum sequence recovery of 53.6%, 60.6%, and 41.5%, respectively, for MHC-I, and 38.3%, 44.6%, and 19.6% for MHC-II. Designed amino acids of three MHC-I test cases from ProteinMPNN, ESM-IF, and Rosetta are represented as sequence logos in Figure 1F. It is not clear whether the difference in MHC-I and MHC-II recovery rates is related to particular structural features or simply a consequence of the reduced number of MHC-II test cases available. Given the more comprehensive number of available 3D structures, we focused the subsequent analysis on only the MHC-I test cases.

Increasing the number of generated designs (up to 500) increased the maximum sequence recovery for some MHC-I test cases (Figure S4), reaching an average maximum sequence recovery of 59.9% (*∼*6% higher than 53.6% with 10 sequences sampled, but not significantly) and 69.1% (9% higher than 60.3% with 10 sequences sampled) for ProteinMPNN and ESM-IF, respectively. Although this increase in sequence recovery was not consistently observed across all test cases (Figure S4), these findings suggest that, at least for some cases, expanded sampling may contribute to the generation of designs closer to the native sequence.

From a practical standpoint, in silico generated designs usually need to be ranked and prioritized for subsequent robust in silico evaluation or experimental validation. Particularly for ProteinMPNN, which provides a score for each design sequence, we tested to rank their designs in the expanded sampling set by the provided ProteinMPNN score (defined as a negative average log-probability) to obtain the top 10 best-scored designs. Although in some cases, such as 8d5q, we observed designs with lower scores (the lower the score, the higher the confidence on the design prediction), achieving high sequence recovery rates, this trend was not consistently observed across all test cases (Figure S5). Additionally, when correlating the scores of all designs with the sequence recovery, we obtained only a moderate correlation (−0.44; Figure S5B). When comparing the sequence recovery of the top 10 ranked designs with the previous 10 designs generated by ProteinMPNN, we did not observe significant differences (average of 44.9% in comparison to the default protocol’s 43.9%) (Figure S6). Together, these findings indicate that, at least for the CDR3 interface design strategy, 10 designs generated by ProteinMPNN would be sufficient to achieve suitable sequence recovery rates.

To investigate the contribution of each designed position to the sequence recovery, we computed the sequence recovery per CDR3 position (Figure 2A). In general, the sequence recovery was uniformly distributed across the CDR3 positions. In CDR3α, position 133, typically located in the middle of the CDR3, exhibits high sequence recovery in both ProteinMPNN and ESM-IF. More pronounced differences between ProteinMPNN and ESM-IF are observed at CDR3α position 134. In CDR3β, ProteinMPNN achieved the highest sequence recovery at position 109, which usually corresponds to the second serine residue of the CASS motif derived from the V gene. This might suggest some bias from model training. In CDR3α, position 114 shows low sequence recovery for ProteinMPNN, contrasting with the recovery observed for ESM-IF. However, a limitation of this analysis is that CDR3 sequence positions can vary substantially in terms of 3D structure positioning. Therefore, we mapped the per-position sequence recovery of ESM-IF onto the TCR structures (Figure 2B). Consistent with the distribution analysis, after TCR superposition, we observed balanced sequence recovery over the TCR interface (Figure 2B, left panel). A possible tendency of higher sequence recovery is noted in a region between CDR3β and CDR3α, whereas CDR3 peripheral regions appear to exhibit lower sequence recovery (Figure 2B, left panel). By superposing the TCR:pMHC structures by the MHC and mapping the sequence recovery onto the CDR3, we also observe evenly distributed recovery, suggesting no bias toward an interacting region of the pMHC (Figure 2B, right panel).

**Figure 2:**
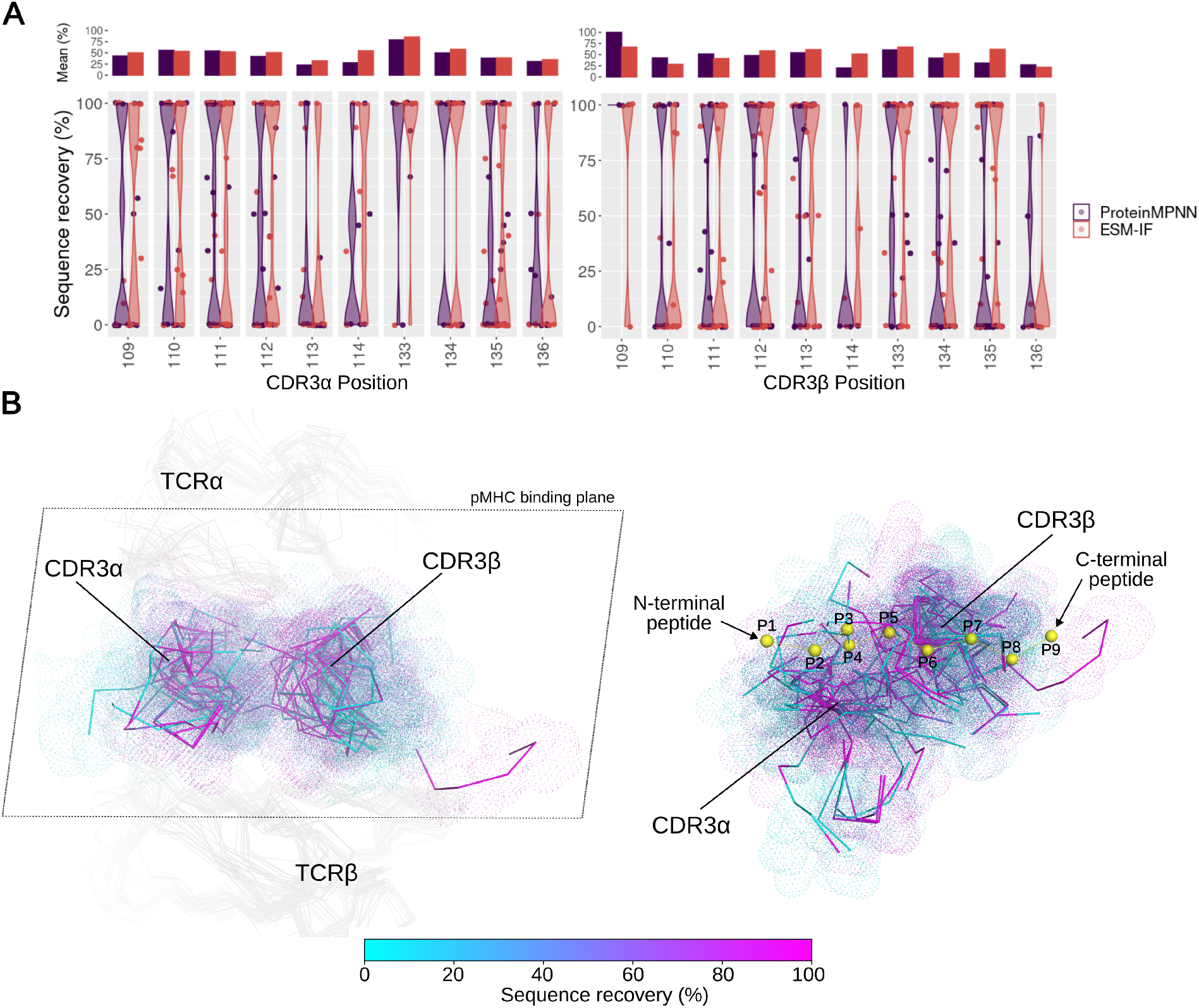
ProteinMPNN and ESM-IF sequence recovery per CDR3 designed positions. **(A)** The distribution of the percentage of sequence recovery per designed CDRα (on the left) and CDR3β (on the right) position is depicted as violin plots. Each point represents the average sequence recovery over non-redundant designed amino acids for a given test case at a given position. The position numbering follows the AHO numbering scheme. The upper bar plot shows the average over each sequence recovery distribution. Positions with fewer than three designed cases were removed for clarity. **(B)** ESM-IF sequence recovery is mapped onto the TCR structures. On the left, structures of test cases are superposed by the TCR, and only the TCR α and β chains are presented. The structures are oriented towards the pMHC plane. On the right, the structures are superposed by the MHC, and only the CDR3s are shown. A representative peptide is presented as yellow spheres to highlight the orientation of the CDR3 segments in relation to the pMHC interface.

Additionally, besides the design of TCR CDR3 interface positions, we explored two other design scenarios: the design of all CDR3 positions (α and β chains), not limited to those at the pMHC interface, and the design of CDR1, CDR2, and CDR3 positions at the interface with the pMHC. Interestingly, when comparing the sequence recovery across the different ProteinMPNN design scenarios, we observed higher recovery rates when designing all CDR3 positions (52.6% compared to 43.9%, on average, over all test cases), rather than only those at the pMHC interface (Figure S7). However, this may be a bias derived from the N and C termini of the CDR3s, which are conserved across different V and J germlines and could potentially be seen during model training. In turn, allowing the design of all the CDRs at the interface with the pMHC reduced the sequence recovery, while increasing diversity (38.3% in comparison to 43.9%, on average, over all test cases) (Figure S7). This can be explained by the fact that in canonical TCR binding modes, the CDR3 are usually buried at the center of the TCR:pMHC interface, whereas the other CDR (CDR1 and CDR2) located at peripheral regions are more prone to exposure to the solvent and consequently at a position less restrictive to different amino acids. A similar trend was noted in the designs generated by ESM-IF. It is noteworthy that ESM-IF utilizes the UniRef sequence dataset for model training [21]. While full CDR3 sequences are absent in TCR sequences from UniRef, as they are generated through gene recombination, other CDRs that constitute part of the TCR V germline could potentially be included in the ESM-IF training set. However, upon analyzing the results from all CDR designs, we did not observe any overestimation of sequence recovery for these cases.

Overall, these findings underscore the capacity of ProteinMPNN and ESM-IF, utilizing fixed backbone positions, in generating TCR amino acids closely resembling native sequences in comparison to Rosetta Design. Given the superior performance of these methods, we confine the subsequent analysis presented in the following sections to deep learning methods. Additionally, to prevent any potential training information leaks to our test cases, we focused the subsequent analyses on the design of TCR CDR3 interface positions. This scenario is also more relevant since CDRs other than CDR3 can establish extensive contacts with the MHC, thereby increasing the risk of cross-reactivity.

### 2.2 Deep learning-based methods recovered the interface amino acid composition for the majority of amino acids

The preservation of interface amino acid composition is crucial, particularly for the TCRs, as it plays a key role in maintaining binding specificity and leads folding into a loop structure. Here, we compared the composition of the CDR3 interface in native sequences and design sequences (Figure 3A).

**Figure 3:**
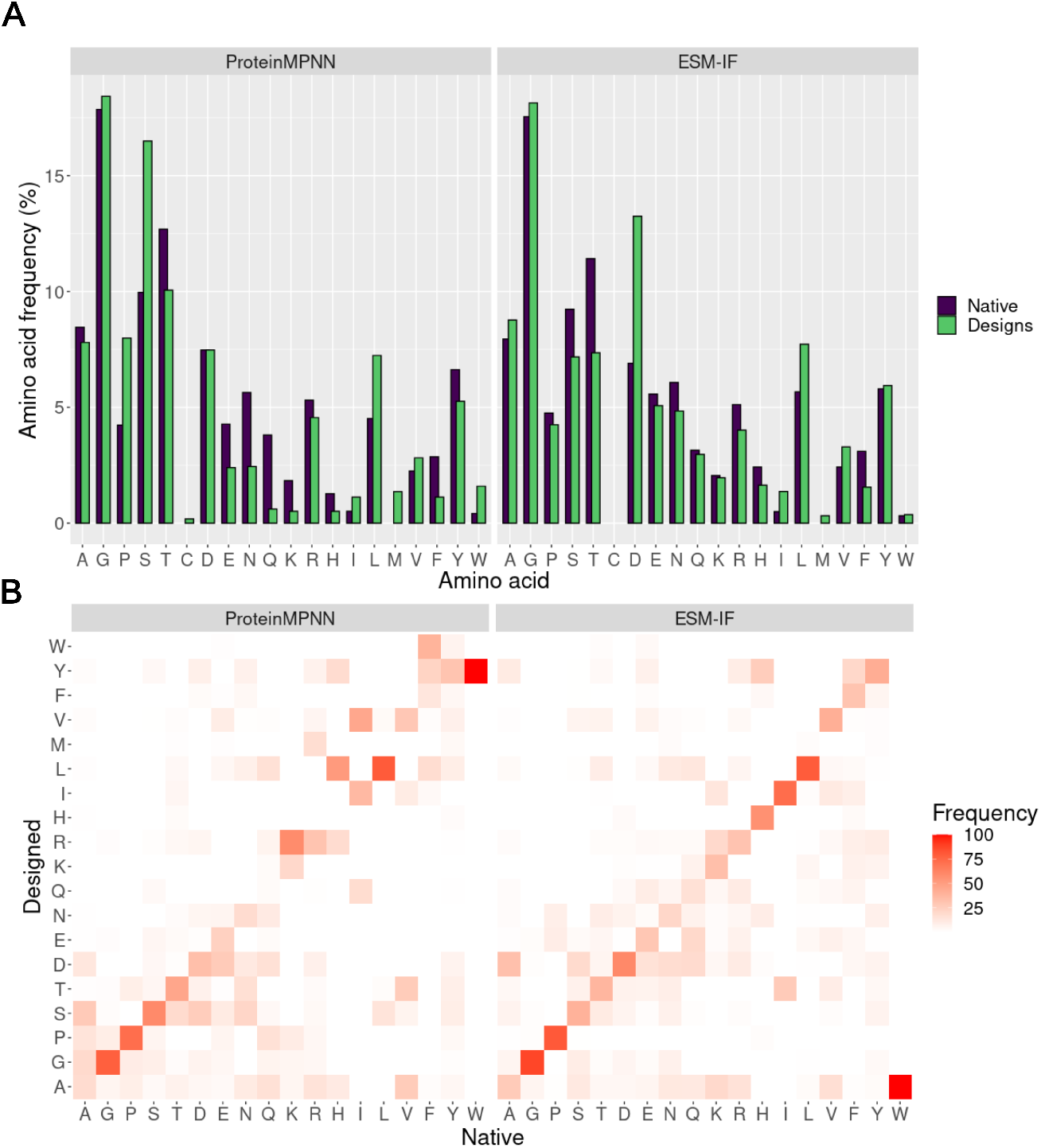
Occurrence of amino acids at CDR3 interface positions in native and designed sequences. **(A)** Frequency of each amino acid at the CDR3 designed positions in native and ProteinMPNN (left panel) or ESM-IF (right panel). Higher frequency indicates that the amino acid was more frequently observed at the CDR3 interface in the analyzed test cases. Only non-redundant generated sequences are considered in the analysis. The x axis is ordered by Dayhoff amino acid grouping: Small (A, G, P, S and T), Sulfur polymerization (C), Acid and amide (D, E, N and Q), Basic (K, R and H), Hydrophobic (I, L, M, V) and Aromatic (F, Y, W). **(B)** Heat map of the frequency of substitutions of the amino acid substitutions in the designed sequences. The x axis represents the amino acids at the native sequences and the y axis represents the corresponding substitution in the designed sequences. A hypothetical frequency of 100% alanine in native sequences and leucine in designed sequence, for instance, indicates that we observed a change from alanine to leucine in all design cases.

Across the CDR3 (α and β chains) interface of all targets, the designed interfaces generated by both ProteinMPNN or ESM-IF generally mirrored the amino acid composition of the native interfaces. As expected, interface CDR3 positions exhibited a strong bias toward glycine, which was recapitulated by both designing methods. Amino acids with low occurrence in native interfaces, such as cysteine, histidine, isoleucine, lysine, methionine, and tryptophan, were also scarce in the design sequences. However, notable differences were observed, including an increased occurrence of leucine, proline, and especially serine in the ProteinMPNN-designed sequences, along with a reduction in acidic and amide amino acid groups (glutamic acid, glutamine, and asparagine). In ProteinMPNN, some native alanine, threonine, and aspartic acid were substituted by serine, leading to an increased frequency of the latter at the CDR3 interface (Figure 3B). Conversely, the reduction in the occurrence of acidic and amine amino acid groups was not observed in ESM-IF which seemed to maintain more balanced substitutions than ProteinMPNN, as expected due to its higher sequence recovery (diagonal trend at the heatmap). Notably, a bias towards aspartic acid was observed in sequences generated by ESM-IF.

### 2.3 Physicochemical related substitutions and sequence recovery at buried or hotspot positions

Since substitutions by physicochemical similar amino acids at protein interfaces can preserve the interface characteristics and, consequently, protein binding, we evaluated the contribution of physicochemical related substitutions to the MHC-I sequence recovery rate by ProteinMPNN and ESM-IF. Approximately 24% and 18% of the CDR3s positions, which underwent amino acids changes in the design process by ProteinMPNN or ESM-IF, respectively, exhibited substitutions to physicochemical related amino acids (according to BLOSUM grouping, see methods for details). Considering this similarity metric, the averaged sequence recovery increased from 43.9% to 56.9% and from 50.1% to 59.0% for ProteinMPNN and ESM-IF, respectively (Figure 4).

**Figure 4:**
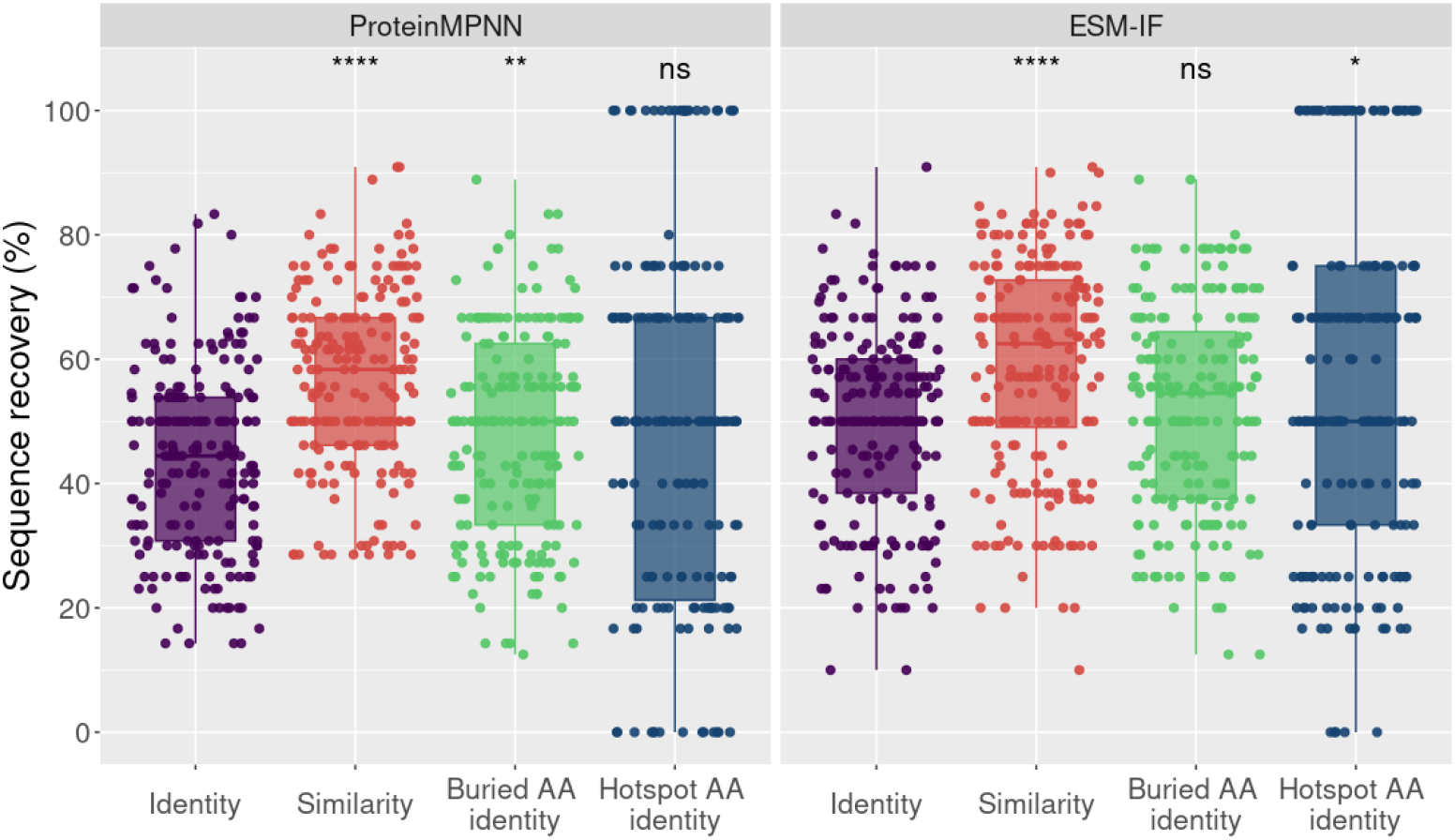
Sequence recovery analysis of interface CDR3s amino acids in terms of identity, similarity, identity at buried positions and identity at hotspot positions for ProteinMPNN (left panel) and ESM-IF (right panel). For both panels, each point of the box plot represents the percentage of sequence recovery of a unique design sequence from the MHC-I test cases, without redundancy. While identity considers only substitutions to the same native amino acid as recovered, the similarity considers as recovered substitutions to the same amino acid (AA) physicochemical class (see methods). The buried AA identity corresponds to the identity only computed over interface CDR3s hotspot positions (averaged SASA < 5Å2, see methods), whereas the hotspot AA identity corresponds to the identity only computed over interface CDR3s hotspot positions, predicted by computational alanine scanning experiments (see methods). Statistical pairwise comparison assessed the significance between the identity (reference) and the other metrics. It was performed using the Mann-Whitney test with the R ggpubr package. Significance is indicated above each box plot (****, ** and * correspond to a p-value below 0.0001, 0.01 and 0.05, respectively, while ‘ns’ means no significance (p-value *≥* 0.05)). A detailed view of the same evaluated metrics per PDB test case is presented in Figure S8.

Another potential factor influencing sequence recovery is that, even within the interface, certain amino acids at specific positions are not entirely buried, thus allowing less restriction in accommodating different amino acids. By focusing exclusively on designed amino acids located at buried interface positions (average SASA *<* 5Å^2^, see methods), we observed a slight but significant increase to 48.1% in the sequence recovery for ProteinMPNN. However, for ESM-IF we did not observe a significant increase in the sequence recovery at buried positions.

Given that even buried positions within the interface may not always be critical for the interaction, we identified positions that could serve as hotspots for pMHC engagement. Through computational alanine scanning, we pinpointed CDR3 positions that were more crucial for the interaction and assessed the sequence recovery at these positions. Despite observing a significant change for ESM-IF, overall, we did not clearly observe an effect on evaluating sequence recovery at hotspots positions.

Taken together, these results suggest that the interface recovery by deep learning methods could potentially be even higher when considering physicochemical related substitutions, without being strongly influenced by the solvent exposure of positions or their significance as hotspots for the interaction.

### 2.4 Designs generated by ProteinMPNN and ESM-IF are dependent on the pMHC interface

Despite the notable sequence recovery achieved in deep learning-based designs, there remains a possibility that the model predominantly learns the probability of an amino acid being part of the CDR3 backbone structure or inter/intra TCR chain contacts rather than specifically interacting with the pMHC. To investigate this, we conducted a similar design protocol but excluded the pMHC, effectively removing it from the bound TCR:pMHC structure (Figure 5).

**Figure 5:**
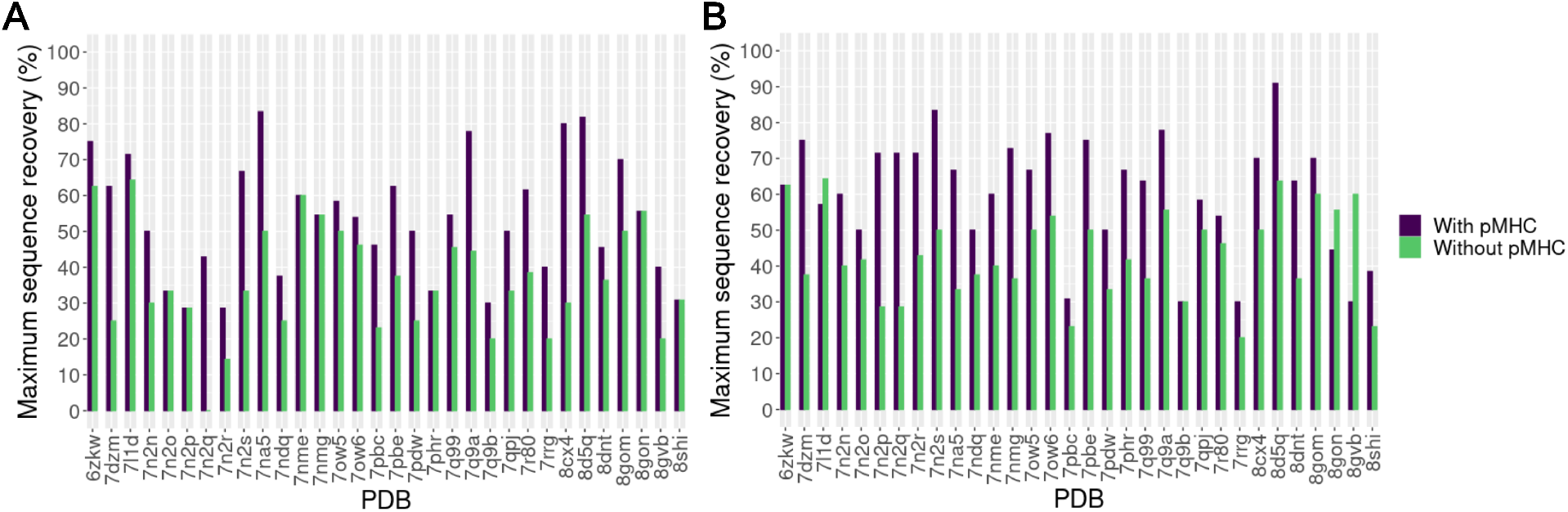
Maximum sequence recovery of interface CDR3s AA with ProteinMPNN (A) and ESM-IF (B) in the presence or absence of the pMHC structures. Bars present the maximum sequence recovery of a design generated by ProteinMPNN (A) or ESM-IF (B) for each test case. Purple bars represent the CDR3 interface designs considering the corresponding pMHC complex whereas the green bars represent the interface design of unbound TCRs, without the pMHC.

As expected, the analysis of maximum sequence recovery revealed a substantial reduction in sequence recovery in 25 (for ProteinMPNN) and 27 (for ESM-IF) out of 32 cases. This indicates that, at least in this analyzed system, the sequence recovery is not solely a consequence of the backbone structure, indicating that the models have learned to design the TCR interface. Intriguingly, in certain cases, such as the 7nme for ProteinMPNN and 6zkw for ESM-IF, the maximum sequence recovery remained unchanged. This observation may suggest that for specific cases, the backbone conformation, or even TCR inter/intra chain contacts, play a crucial role in the design process.

### 2.5 AlphaFold2-based modeling supports the ProteinMPNN and ESM-IF TCR designs

In addition to the design evaluation at sequence-level, we performed a structural analysis of TCR designs using a deep learning-based approach. We employed TCRModel2, an AlphaFold2 modified version focused on TCR modeling [25]. Design models exhibiting low atomic deviation from the native structure and high modeling confidence score [25] (Figure S9) indicate that TCRModel2 modeling supports the interface designed by ProteinMPNN or ESM-IF, reinforcing our confidence in the generated design. It is important to note that we restricted our analysis to cases where native TCR:pMHC structures from the MHC-I test cases were accurately remodeled with medium or high accuracy (77% of cases) according to CAPRI criteria assessed by DockQ scores. Furthermore, an additional filter was applied based on the TCRModel2 confidence score (*≥* 0.85). When combined, these criteria selected 60% of the cases for use in the structural analysis of designs.

In 84% and 89% of the analyzed cases, respectively, the ProteinMPNN or ESM-IF design with the highest model confidence reached similar or better model confidence than the median of remodeled native sequences. In seven and eight cases, for ProteinMPNN and ESM-IF, respectively, the designs achieved a model confidence higher than 0.9 (Figure 6A). When randomly selecting dissimilar amino acids (see methods) at the same positions (excluding cysteine) instead of generating amino acids by ProteinMPNN or ESM-IF, we can see a clear drop in the model confidence and we observed random designs with the highest model confidence reaching similar or better model confidence than the median of remodeled native sequences in only 48% of the cases (Figure S10A). Additionally, we also observed a higher variability in the model confidence of random designs, with some designs reaching confidence scores below 0.6. It is worth noting that TCR:pMHC interfaces are relatively large and do not involve only CDR3 contacts, thus the occurrence of random designs with high confidence is not unexpected.

**Figure 6:**
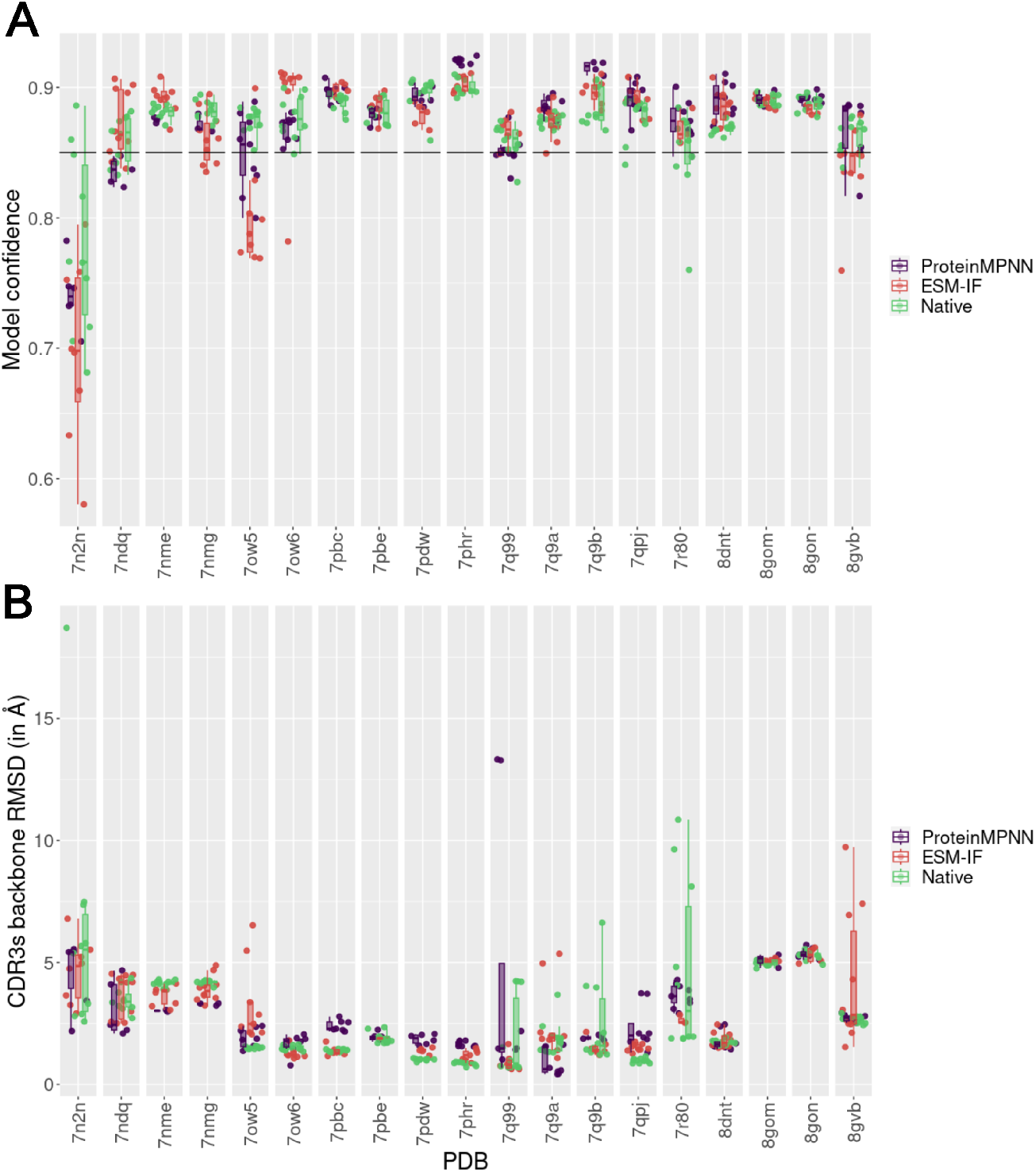
Modeling of TCR designs with TCRmodel2. **(A)** Box plot of model confidence of ProteinMPNN (in purple) and ESM-IF (in red) TCR designs for each test case. Each point represents a different design. The model confidence of remodeled native sequences is colored in green. In this case we remodeled the native sequences 10 times to obtain a distribution of modeling scores that represent the native structure. **(B)** Root-mean-square deviation (RMSD) of CDR3 backbone atoms (both alpha and beta TCR chains) of designs in comparison to the corresponding native crystal structure that originated the designs. RMSDs were determined after structure superposition by the MHC.

We also assessed the deviation between the CDR3 backbone in the designs and the native structure, comparing these deviations to those observed in the remodeled native sequence. Even minimal backbone deviation could be expected due to side chain readjustments required to accommodate designed residues. In 79% of cases for ProteinMPN and 89% for ESM-IF, at least one design achieved a comparable or lower CDR3 backbone RMSD compared to the median of remodeled native sequence (Figure 6B). Deviations below 2 Å were observed in 53% of cases for both deep learning models. When considering both model confidence and RMSD, in most cases, we observed that the designs, alongside the native sequences, exhibited optimized model confidence and RMSD (Figure S10C).

Collectively, these findings demonstrate that in the majority of tested cases, the designs generated by ProteinMPNN or ESM-IF were successfully modeled by TCRModel2 with high confidence scores, showing relatively low deviation from the CDR3 native structure. These results suggest that AlphaFold2-based modeling supports the potential binding of most TCR designs to the pMHCs in the test cases.

### 2.6 Energetic scoring of the TCR designs

As another structure-based criterion to evaluate the TCR designs, we employed the Rosetta energy function to score the designed structures against the corresponding native TCR complexes (the same set used in the AlphaFold modeling assessment). The scoring was based on two widely used Rosetta terms obtained from the Rosetta InterfaceAnalyzer: *dG_separated* and *total_score*. While the former relies on predicted energy changes at the binding interface, the latter provides broader information about complex stability. These two terms can be combined in the design evaluation [26].

The *dG_separated* scoring indicated that both ProteinMPNN and ESM-IF designs followed a similar scoring profile to that determined for the native structures in most test cases (Figure S11A). Designs with worse score, i.e., higher *dG_separated* scores, and higher score difference than native structures were observed in only two test cases (7pdw and 8dnt) for ProteinMPNN. Conversely, ESM-IF showed a design with equal or better *dG_separated* score, than the native structure in all test cases. This contrasts with the scores obtained from dissimilar random TCR sequences, which presented higher variability and worse score and higher score differences than native structures in five test cases. This behavior is more pronounced when considering complex stability through the Rosetta *total_score* (Figure S11B). ProteinMPNN and ESM-IF designs clearly followed the energetic profile of native structures in all test cases, contrasting with the worse scores, i.e., higher *total_score*, presented by the dissimilar random TCRs. A combination between the two Rosetta terms can be observed in Figure S11C.

### 2.7 Molecular dynamics simulations and computational binding affinity estimation of the TCR designs

To assess the binding affinity of the TCR designs and compare them with native TCRs, we utilized a robust molecular dynamics simulation protocol coupled with widely used free energy estimation based on a previously benchmarked strategy relying on MM/PBSA calculations [27].

While this MM/PBSA protocol has shown promising results for estimating the binding free energy of two specific TCRs (1G4 and A6), its capacity to generalize to other distinct TCRs remains unexplored. Hence, as a first step to ensure a reliable evaluation of the binding affinity of our TCR designs, we benchmarked this protocol to predict free energy changes resulting from mutations in a curated set of TCR:pMHC wild-type and mutated paired complexes, all possessing solved 3D structures and experimental binding affinity determinations. This benchmark test set comprised 5 different TCRs, with mutations ranging from single-point to 12 mutations across TCR α and β chains. By applying this MM/PBSA protocol, we achieved a notable correlation coefficient of 0.9 (Pearson correlation) with the experimental data (Figure S12), thereby confirming its efficacy for comparing the binding affinity of native and TCR designs.

Since this MD-based protocol is time-consuming, we selected designs from 7 TCR:pMHC test cases to perform the MM/PBSA calculations over MD simulation trajectories (Figure 7). The trajectories of the designs were stable, maintaining CDR3 backbone deviation medians below 2.5 Å, in most test cases, similarly as the native trajectory (Figure S13). In general, the Principal Component Analysis (PCA) indicated that the designed CDR3s sampled a similar conformational space as the native ones, with some exceptions like designs from the 7n2o test case (Figure S14). However, the deviation between these different sampling spaces is low (below 1 Å), as indicated by the RMSD medians (Figure S13).

**Figure 7:**
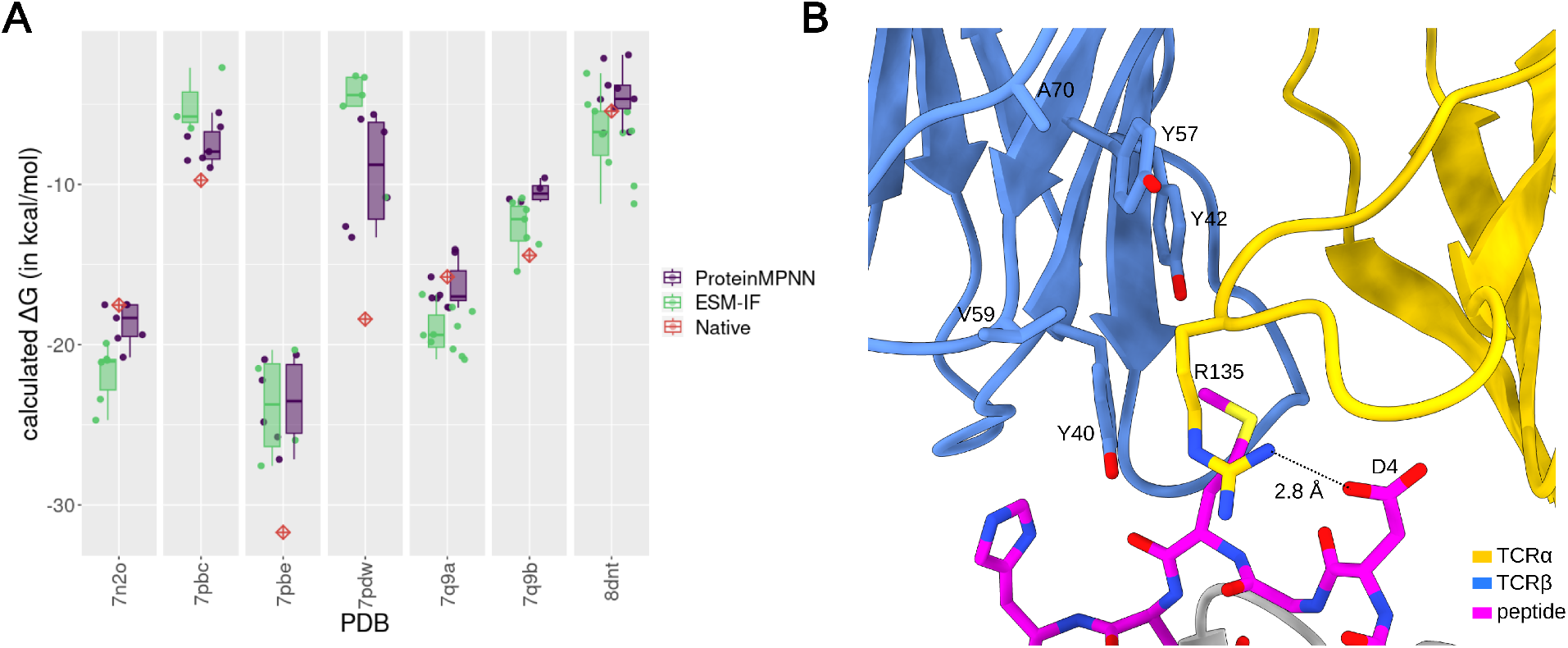
Molecular dynamics simulation analysis and binding affinity estimation with MM/PBSA of Native and ProteinMPNN and ESM-IF TCR designs. **(A)** Box plot of calculated ΔG (in kcal/mol) of ProteinMPNN (in purple) and ESM-IF (in green) designs in comparison to the calculated ΔG of the native complex (red diamond) for each evaluated MHC-I test case. Each point corresponds to the median of calculated ΔG from 15 replicas. The detailed distribution of ΔG across replicas and statistical tests are presented in Figure S15. **(B)** Visualization of the TCR and peptide interface from the PDB 7pdw, highlighting the interaction between R135 from the TCRα and the D4 residue of the peptide.

Interestingly, in most of the test cases (5 out of 7), we observed designs presenting binding affinity (expressed by ΔG) similar to or better than their corresponding native structure (Figure 7A and Figure S15). Most designs, whether from ProteinMPNN or ESM-IF, recovered the native binding affinity, presenting no significant difference in comparison to the native ΔG. Nonetheless, in one test case (7q9a) for ProteinMPNN and in three test cases for ESM-IF (7n2o, 7q9a and 8dnt), we obtained TCR design with significant higher affinity (lower ΔG) than the native TCR (assessed by a Mann-Whitney test, see Figure S15). Intriguingly, both methods failed to generate designs with compatible affinity to the native TCR for 7pbe and 7pdw test cases.

A free energy decomposition of the native 7pdw interaction revealed that most of the TCR interaction relies on TCR three arginine from the TCRα (Figure S16). Particularly, the one with most contribution (R135) anchors to the D4 peptide using a rotamer conformation that allows this interaction even with a high Cα distance and Cβ atoms pointing in opposite directions. The proposed amino acid by ProteinMPNN is a methionine which could interact with hydrophobic residues of TCRβ like V59, while the proposed amino acids by ESM-IF are the glutamine or glutamic acid, which could create a polar interaction and/or hydrogen bondings with a cluster of tyrosine residues of TCRβ (Figure 7B). Consequently, the positioning of this arginine hotspot together with the chemical environment can pose a strong challenge to these deep learning methods based on fixed backbones.

## 3 Discussion

In this study, we investigated the utilization of two recently developed deep learning-based protein design methods, ProteinMPNN and ESM-IF, for designing TCRs interfaces. The primary aim was to evaluate the capabilities of these methods in designing TCRs capable of binding to a specific antigenic peptide presented by the MHC, thereby guiding computational design strategies and impacting on TCR engineering.

Despite the advances in protein language models, the limited quantity and quality of TCR sequence data with known specificity to antigens hinder the direct use of generative models only based on amino acid sequences [28]. As a result, an alternative approach to tackle the computational design of TCRs is to base the design process on the 3D structure of the complex formed by the TCR and the peptide presented by the MHC. Indeed, structure-based design of TCRs is a traditional approach that, thus far, relies solely on parameterized energy functions to predict amino acid substitutions that could enhance TCR interactions [14, 18, 19].

Unlike other immunological interactions, such as those between antibodies and antigens, TCR interactions possess several distinctive characteristics. One of the primary distinctions is the binding affinity, which typically falls within the micromolar range. The therapeutic use of TCRs does not necessarily require optimized affinity, since supraphysiological affinities can even prove detrimental to T cell response [14, 29, 30]. These unique characteristics of TCR interaction underscore the need to explore alternative methods for computational design, strategies that not only rely on numerical energy functions but also have the capability to capture native features of this interaction.

In this study, we designed TCRs to bind antigenic peptides presented by MHCs based on an initial 3D structure of the complex using ProteinMPNN and ESM-IF with fixed-backbones. To assess the quality and success of these designs, we compared them to experimentally determined TCRs with solved 3D structures bound to the pMHC, serving as the initial models for the design process. It is worth noting that structure-based design approaches are particularly useful when aiming to enhance an existing interaction or even modify it to minimize cross-reactivity. However, recent advances in generative protein backbone methods, such as RFDiffusion [31], have overcome the limitation imposed by the need for experimentally determined initial structure models, thereby enabling the generation of protein backbones to target a specific protein, which can subsequently be designed using deep learning methods.

The evaluation of designs was based on a comprehensive set of orthogonal computational approaches, primarily utilizing two different kinds of metrics: sequence-based and structure-based metrics. At the sequence-based level, we found that the deep learning methods were able to recover a substantial percentage of native amino acids at the designed positions, outperforming a physics-based method like Rosetta Design. Notably, when comparing ProteinMPNN and ESM-IF, we took care to conduct a detailed temperature sampling analysis to ensure comparability for this specific task. Interestingly, in our analyzed test cases, the sequence recovery of ESM-IF was generally higher than that of ProteinMPNN. Despite both methods being based on graph neural networks, the training sets differ significantly. While ProteinMPNN is tranied on non-redundant solved 3D structures available in the PDB, comprising approximately 25,000 clusters of heteroligomeric assemblies, ESM-IF was trained on single-chain 3D structures composed by the combination of 12 million modeled UniRef sequences and 16,000 solved structures. Despite not being explicitly trained on oligomeric interfaces, ESM-IF was able to recover native sequences, even outperforming ProteinMPNN, which could be attributed to the larger number of training samples.

Of note, the sequence recovery obtained in our tests is comparable to that reported in the original proteinMPNN study (51% of recovery for heteromeric interfaces) and ESM study as well [20, 21]. This indicates that despite the peculiarities of TCR interfaces, such as being composed of loops, they did not affect the capability of those methods in recovering the native interface. Specific biases observed for general ProteinMPNN designs [20], such as reduction in glutamine incorporation, are also evident in our ProteinMPNN designs. However, in our ProteinMPNN designs we observed a reduced presence of charged amino acids, particularly glutamic acid and lysine, and an increased presence of serine, which contradicts a tendency observed in ProteinMPNN general designs to convert polar residues into charged amino acids [20]. We hypothesize that these differences may be a consequence of the unique characteristics of the TCR:pMHC interface.

Importantly, the estimated sequence recovery was associated with the presence of the pMHC, suggesting a binding-dependent nature, as the same design in the absence of the target pMHC substantially reduced the recovery of native amino acids. This is significant to demonstrate that the predictions, even from a model such as ESM-IF that was only trained on single-chain structures, are not solely based on the TCR backbones or even TCR intrachain contacts but are specifically dependent on the target pMHC.

Despite sequence recovery being a valuable and widely used metric for assessing the performance of protein design methods, TCRs with low sequence recovery can still exhibit binding. Indeed, diversity is usually advantageous in a protein design context, particularly for TCR:pMHC interfaces, which may lack naturally optimal interactions [8]. Additionally, it is possible that even with high sequence recovery, a single amino acid divergence compared to the native sequence can have significant effect on the protein stabilization and binding. For this reason, we evaluated the quality of designs using a combination of structure-based metrics. Both AlphaFold2-based modeling and Rosetta scores suggested that the TCR designs generated by ProteinMPNN or ESM-IF can form stable complexes with target pMHCs at a similar level to native TCRs.

As a critical characteristic of TCR interfaces, we assessed the binding affinity of the designed TCRs to pMHC targets using a benchmarked protocol of MD simulations with MM/PBSA for the estimation of the ΔG of binding. Our results suggested that, for most cases, the TCR designs, whether from ProteinMPNN or ESM-IF, recapitulate the binding affinity of the native TCR. In only two test cases did the models fail to generate TCRs with compatible affinity to the native TCRs. These cases, such as the 7pdw case, may serve as study cases for improving the design methods and highlighting the limitations of backbone design methods. Although ESM-IF was able to generate enhanced TCRs (as observed in the 8dnt test case), the improved ΔG was not substantial, emphasizing that these methods alone may not consistently optimize the interface. If an optimized interface is required, a combination with an optimization algorithm, as reported in [23], could be beneficial.

A critical limitation posing challenges to the practical applicability of engineered TCRs is their potential for cross-reactivity, exemplified by the fatal cross-reactivity observed in a clinical study with an affinity-matured TCR that was reactive against the human titin self-peptide [15]. Although cross-reactivity is not the focus of this study and therefore not assessed for the deep learning-generated TCR designs, we believe that the advancements in these protein design methods can contribute to the development of more specific TCRs. These can be further investigated by cross-reactivity predictive methods [32, 33].

In conclusion, our study demonstrates that deep learning design methods such as ProteinMPNN and ESM-IF, which utilize fixed-backbone 3D structures of TCR:pMHC complexes, can propose amino acids at the TCR interface with high similarity to native sequences, predominantly guided by pMHC contacts. Computational structure-based metrics indicate that the designed TCR complexes are stable, exhibiting a binding affinity similar to native TCRs, with some cases even showing improved affinity. These findings contribute to advancements in the field of TCR design and, consequently, in T cell-based immunotherapy.

## 4 Methods

### 4.1 Benchmarking dataset

To assess the capability of computational design methods for TCR designing, we curated a dataset of TCR:pMHC complexes, involving both MHC-1 and MHC-II, with solved 3D structures that are not included in the ProteinMPNN training dataset (August 31, 2021, date cut-off) and not in AlphaFold2.3 (September 30, 2021, date cut-off). This date cut-off ensures that the complexes under consideration were not part of ProteinMPNN or AlphaFold2.3 training datasets, which could potentially lead to an overestimation of sequence recovery or modeling accuracy results. ESM-IF training involves the utilization of protein models derived from UniRef sequences and experimentally determined structures from the CATH 4.3 database. However, it’s important to note that the CDR3 sequences, which are central to this study, are not included in the UniRef dataset as they are generated through gene rearrangement. Additionally, the CATH 4.3 release occurred in 2019, which falls within the data cut-off period for ProteinMPNN and AlphaFold2.3.

The structures were collected from the TCR3d database [34] on December 13, 2023, and selected based on a resolution cut-off of 3.25 Å. We verified whether the chosen TCR:pMHC complexes had a related PDB entry released before the cut-off dates. For this reason, the CDR3β sequences of the TCR from each complex were submitted to a blast search using the blastp server [35]. Additionally, we verified redundancies within the selected dataset of TCR:pMHC complexes (considering the same TCR, peptide and MHC). For this reason, complexes with different peptide or MHC sequences, despite having the same CDR3, were maintained in the dataset (Table S1). Structures with unnatural amino acids were excluded from the dataset. Following these filtering steps, a total of 32 and 6 TCR:pMHC structures (MHC-I and MHC-II, respectively) were selected, as listed in Table S1.

### 4.2 ProteinMPNN and ESM-IF design protocols

For ProteinMPNN design, we used the v_48_020 model with default backbone noise (0.00) and sampling temperature of 0.1 to generate 10 designs per each test case, unless otherwise specified. To evaluate the effects of sampling temperature on sequence generation, we utilized a range of temperatures from T=0.000001 to 5. Additionally, to assess the impact of the number of generated sequences, we varied the quantity of generated sequences per test case from 5 to 500. Three design strategies were evaluated: (1) restricted the design to CDR3 (α and β TCR chains) within a proximity of 5 Å to either the peptide or MHC, (2) CDR1, CDR2, or CDR3 positions (α and β TCR chains) within a 5 Å distance to the peptide or MHC, or (3) the entire CDR3 (α and β TCR chains). Positions outside the specified criteria were held constant, and for the designed positions, all amino acid substitutions, including cysteine, were allowed. All tests were performed maintaining the same seed.

For ESM-IF design, we employed the esm_if1_gvp4_t16_142M_UR50 (fair-esm v2.0.1) model with sampling temperature of 0.2 to generate 10 designs per each test case, unless otherwise specified. Similarly to ProteinMPNN, effects of sampling temperature on sequence generation were evaluated using a range of temperatures from T=0.000001 to 5, and the quantity of generated sequences per test case varied from 5 to 500 to assess the impact of the number of generated sequences. Three design strategies were explored: (1) restricting the design to CDR3 (α and β TCR chains) within a proximity of 5 Å to either the peptide or MHC, (2) including CDR1, CDR2, or CDR3 positions (α and β TCR chains) within a 5 Å distance to the peptide or MHC, and (3) encompassing the entire CDR3 (α and β TCR chains). Positions outside the specified criteria were held constant. The designed positions were changed to <mask> tokens, and all amino acid substitutions, including cysteine, were allowed. Given the multi-chain structure, a padding of 10 <pad> tokens was used to separate the chains. The ESM-IF design protocol used in our study is available at https://github.com/LBC-LNBio/ESMIFDesign.

### 4.3 Rosetta design protocol

For Rosetta design, we utilized the Rosetta version 3.5.1, and the protocol was executed using RosettaScripts. The target design positions were the same as those in deep learning-based methods, and we utilized a Rosetta resfile to indicate the positions and allowed substitutions (in this case we excluded cysteine) (resfile example in Supplementary Information C.1). Throughout the mutation process, we applied the InterfaceDesign2019 fast relax protocol. A total of 10 designs were generated per test case. Alternatively, we generated a total of 1000 designs per test case and ranked the designs based on the Interface Analyzer *dG_separated* term or the *total_score* term, considering the ref2015 score function to identify the top 10 best-scored designs. A representative example of the RosettaScripts xml file is provided in Supplementary Information C.2.

### 4.4 Design quality assessment at sequence level

To evaluate the success of designs, we compared the designed sequence to the native sequence by using the sequence recovery metric. The sequence recovery (Equation 1) measures the percentage of positions in designed sequences that match the amino acids in the native sequence. Given two sets of corresponding positions *X* = {*x*_1_, *x*_2_, …, *x*_*n*_} and *Y* = {*y*_1_, *y*_2_, …, *y*_*n*_} of equal length *n*, the sequence recovery is defined as:

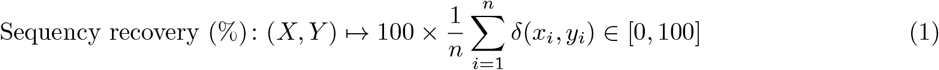

where *δ*(*x*_*i*_, *y*_*i*_) is defined as:

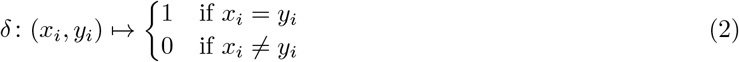

The percentage sequence recovery of 100% indicates that the designed sequence is identical to the native sequence, and 0% indicates no recovery. When evaluating the sequence recovery, we only considered non-redundant sequences with each test case.

When comparing ProteinMPNN and ESM-IF across different sampling temperatures, we employed two additional sequence metrics: Uniqueness and Entropy. The Uniqueness (Equation 3) represents the percentage of unique sequences within the designed set. Given a set of *m* designed sequences 𝒟 = {*D*_1_, *D*_2_, …, *D*_*m*_}, with *u* denoting the count of unique sequences, the Uniqueness is calculated as:

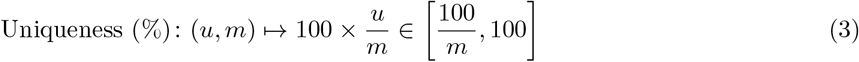

The Entropy relies on Shannon’s information-theoretic entropy (Equation 4) and measures the diversity of the designed sequences. For each position *j* in the designed sequences of length *L*, the probability distribution of amino acids is estimated from the observed frequency. The Shannon’s information-theoretic entropy of the position *j* in the designed sequences is defined as:

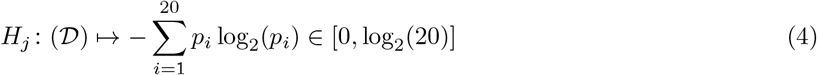

where *p*_*i*_ is the probability of amino acid *i* at position *j* in the designed sequences.

The entropy of each position is then averaged across all positions to obtain the Mean Sequence Entropy (MSE) as defined in Equation 5:

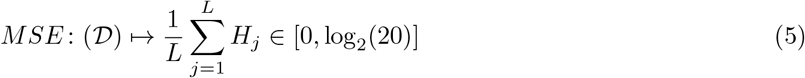

Additionally, we explored physicochemical similarity by considering changes from an amino acid in the native sequence to another in the design sequence within the same group. Amino acid groups were created using hierarchical clustering of BLOSUM62 amino acid features into 8 groups: [A, G, and S], [R, N, Q, H, and K], [D and E], [C], [I, L, M, and V], and [F, W, and Y].

### 4.5 Identification of TCR buried and hotspot positions

To compute sequence recovery over buried positions in the TCR:pMHC complexes of the test cases, we identified buried positions in native structures based on solvent-accessible surface area (SASA) determined for each position using FreeSASA in the vanddraabe R package. We considered a position exposed if the SASA, computed through all atom averaging, was higher than 5 Å.

The hotspot positions were determined by performing an alanine scanning calculation using Rosetta 2.3 software. In this analysis, a residue was considered a hotspot if the absolute calculated ΔΔG for alanine substitution was greater than 0.5 kcal/mol. The alanine scanning command is provided in Supplementary Information C.3, along with an example of the mutation list file used in this protocol (Supplementary Information C.4).

### 4.6 Design quality at structural level

To evaluate the structural quality of ProteinMPNN or ESM-IF TCR designs, we modeled these designs and employed various orthogonal approaches for assessment. This included AlphaFold2-based modeling, Rosetta-based scores and molecular dynamics simulations with MM/PBSA calculations. Further details regarding the latter are provided in the following section.

The ProteinMPNN or ESM-IF design sequences were modeled with TCRModel2, a modified version of AlphaFold2 focused on TCR modeling [25]. The standalone version of TCRModel2, available at https://github.com/piercelab/tcrmodel2/, was employed with default parameters, except for the *max_template_date* flag, which was set to 2021-09-30 (AlphaFold2.3 cut-off date). Relaxation with Amber was allowed. The confidence score was extracted from the TCRModel2 output statistics file. The RMSD between the CDR3 of the design structure and the native structure was estimated considering the CDR3 backbone atoms after superposition of the design structure to the native structure by the MHC. RMSD calculations were performed in R using the Bio3D package [36] and the DockQ calculation was performed using its standalone version obtained from https://github.com/bjornwallner/DockQ [37].

For the assessment of the TCR:pMHC interface and complex quality, we employed the Rosetta Interface Analyzer, utilizing Rosetta software (version 3.5.1), with the entire protocol executed through RosettaScripts. Upon mutations of the native structures according to the designed amino acids, the complex was relaxed with the FastRelax protocol using backbone constraints to prevent structure displacement. The REF2015 score function was applied throughout the protocol, with *dG_separated* and *total_score* terms extracted from the output file. An example of the RosettaScripts used for this purpose is provided in Supplementary Information C.5.

Random dissimilar TCR sequences were generated by substituting the native amino acids at the designed positions with randomly selected amino acids from different groups. Amino acid groups were defined by clustering the amino acids into six groups based on hierarchical clustering of BLOSUM62 features.

### 4.7 TCR:pMHC binding affinity dataset and benchmark

To validate the capability of the binding affinity methods employed in this study to accurately predict TCR:pMHC interaction and stability, we created a dataset of TCR:pMHC complexes with solved 3D structure and experimental binding affinity information. Since our aim in this study is to evaluate TCR designs, we focused the analyses on the paired comparison between wild-type TCR:pMHC complexes and its corresponding mutant. For this reason, we only considered wild-type complexes that have a corresponding TCR mutated sequence with known complex 3D structure.

We selected wild-type and mutant pairs from the ATLAS server [38] and excluded from the dataset samples in which the experimental binding affinity of the wild-type and mutant complex were not obtained from the same study. By doing this, we seek to minimize noise included from different methods or experimental setups. Following these criteria, 12 TCR:pMHC complexes were found, forming 7 wild-type and mutant pairs (Table S2). Since native and mutant pairs could have different sequence lengths as a consequence of N or C terminal electron density confidence, we trimmed the longest sequence in the pair to match the length of the shorter sequence. This was done in order to avoid bias created by the differing number of residues in MD simulations and MM/PBSA calculations.

### 4.8 Molecular dynamics simulations

To estimate the binding affinity of the TCR:pMHC complexes, we first sampled the conformational space of the complexes with molecular dynamics (MD) simulations.

For MD simulations of native and designed TCRs in complex with the pMHC, we obtained the native TCR:pMHC complexes from the Protein Data Bank (PDB). The complexes were composed by the variable and constant TCR domains, including both α and β chains, the peptide, the MHC and the β2-microglobulin. None of the selected complexes had gaps in the structure. The native complex was processed by removing water and small molecules. Protonation state of residues was determined at a pH of 7.0 using Propka 3.0 [39] with pdb2pqr [40]. The designed TCR complexes were built using the native structures as reference and substituting the designed residues with PyMOL Python package. The protonation states of histidine at non-designed positions in designed complexes were maintained the same as the one in the corresponding native complex.

Prior to protein solvation, water molecules were placed on water sites by following a protocol similar to [27]. Specifically, the three-dimensional reference interaction site model (3D-RISM) [41, 42] and the MoFT suite [43] were used to calculate the solvent density around the protein and to find and place discrete water molecules into the most favorable locations, respectively. This was followed by aligning the complexes with its principal axis of inertia to the z axis, in order to reduce the number of waters using cpptraj from AmberTools23 [44]. All structures were solvated in a rectangular water box such that water molecules encompass the box boundary within 14 Å of the protein and were subsequently neutralized by the addition of Na^+^ ions. The Amber ff14SB force field [45] and the TIP3P water model [46] were used for protein and water, respectively. Disulfide bonds were manually bonded using the information derived from the pdb4amber module from AmberTools23 [44]. The topology and initial coordinates for each solvated system were done using TLeap from AmberTools23 [44].

MD simulations were performed using a graphics processing unit (GPU) accelerated Amber from AMBER22 [47, 48]. Each system was subjected to minimization, equilibration, and production following the protocol reported in [27]. Prior to replicas production, a minimization followed by a short MD simulation was conducted. This involved heating the system from 50 K to 298 K and restraining the protein atom positions. Subsequently followed, the entire system is minimized without any restrictions. The process of generating replicas per complex involved seven steps: (1) the system was heated from 25 K to 298 K for 20 ps at NVT with restraints on the CA atoms with a force constant of 5 kcal/mol/Å^2^, (2-5) four NPT simulations of 10 ps, restraining the CA atoms with a decreasing force constant from 4 kcal/mol/Å^2^ to 1 kcal/mol/Å^2^, (6) a 1 ns NPT without any restraint, and (7) a 3 ns NPT production run at 298 K. A total of 15 replicas were run for each system. The production simulations were used for MMPBSA calculations, as well as PCA and RMSD analysis. The temperature was controlled with a Langevin thermostat [49] using a collision frequency of 1 ps^*−*1^, while the pressure was set to 1 bar using a Berendsen thermostat [50] with a relaxation time of 1 ps.

The MD simulations of the benchmarked complexes described in Table S2 followed the same protocol used above for the native complexes. In this case, no mutation needed to be performed since both native and mutant TCR:pMHC complexes were available.

MD simulations of both designed and native TCRs in complex with pMHC were analyzed by computing the root-mean-square deviation (RMSD) of the CDR3 backbone atoms (α and β chains, considering only a range of positions from the first designed positions to the last designed positions) between trajectory frames from production runs and the corresponding native crystal structure. This involved superposing the trajectory frames by the MHC of the corresponding native crystal structure. Both superposition and RMSD calculations were conducted using the R Bio3D package [36] in RStudio.

To assess the conformational space sampled by the designs compared to the native TCRs, we employed principal component analysis (PCA). The PCA was carried out using the pca.xyz function of the R Bio3D package [36] in RStudio, utilizing the coordinates of the CDR3 positions from the superposed trajectory used in RMSD calculations. All systems, their respective replicas and designs were included in the calculation. The first two principal components were utilized in the evaluation of conformational space sampling.

### 4.9 Free energy calculations with MM/PBSA

The MM/PBSA calculations were performed over the 15 MD simulations replicas of 3 ns saving every 10 ps, giving a total of 300 frames each. For each system, a total of 4500 frames were used for the calculation. The ionic strength and the internal dielectric constant were set to 0.15 M and 6, respectively. All other options were set to default. The MPI version of the MMPBSA.py [51] from AmberTools23 [44] was used for free energy calculations through MM/PBSA approximation. Unlike [27], no explicit waters at the interface nor entropy corrections were included in the MM/PBSA calculations.

## 5 Acknowledgments

We are grateful to the Brazilian Biosciences National Laboratory (LNBio), part of the Brazilian Center for Research in Energy and Materials (CNPEM) for accessibility to the High-Performance Computing Cluster and scientific infrastructure. We thank the ProteinMPNN, ESM-IF and Rosetta team for sharing their algorithm and code.

## 6 Funding

This work was supported by the grant #2022/04260-6, São Paulo Research Foundation (FAPESP) (to H.V.R.F) and financed in part by the Coordenação de Aperfeiçoamento de Pessoal de Nível Superior - Brasil (CAPES) – Finance Code 001 [Grant Number 88887.928702/2023-00] (to J.V.S.G) and the National Institutes of Health Grant NIH GM144083 (to B.G.P). The opinions, hypotheses, and conclusions or recommendations expressed in this material are the responsibility of the author and do not necessarily reflect the views of FAPESP.

## A Supplementary Figures

**Figure S1:**
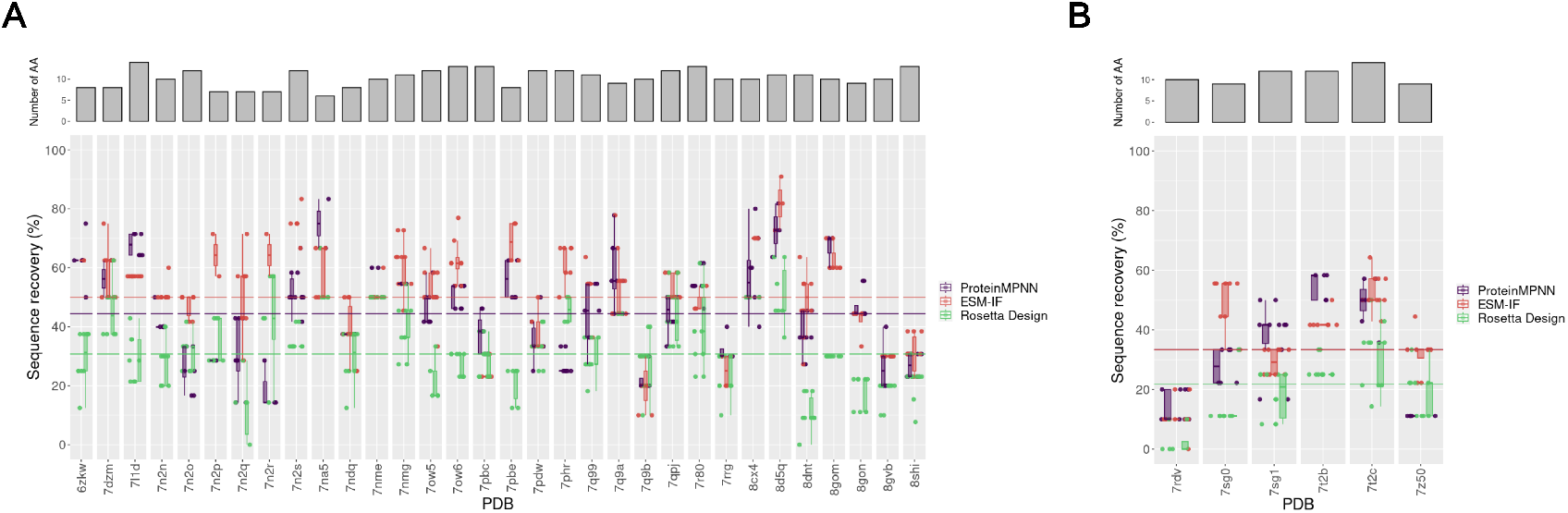
Sequence recovery analysis of interface CDR3 amino acids in designing with ProteinMPNN ESM-IF or Rosetta Design (InterfaceDesign2019 protocol). **(A)** Upper panel shows the number of interface amino acids selected to be designed for each test case. Bottom panel presents boxplots of sequence recovery for each case designed by ProteinMPNN (purple), ESM-IF (orange) or Rosetta (green). Each point corresponds to a design sequence. Redundant designs were removed from the analysis. For both methods, 10 designs were generated per test cases and for Rosetta. Lines indicate the median computed over all designs: 44.4%, 50.0% and 30.8% for ProteinMPNN, ESM-IF, and Rosetta Design, respectively. **(B)** Same as (A), but for MHC-II. Lines are median over all designs: 33.3%, 33.3%, and 21.8% for ProteinMPNN, ESM-IF and Rosetta Design, respectively.

**Figure S2:**
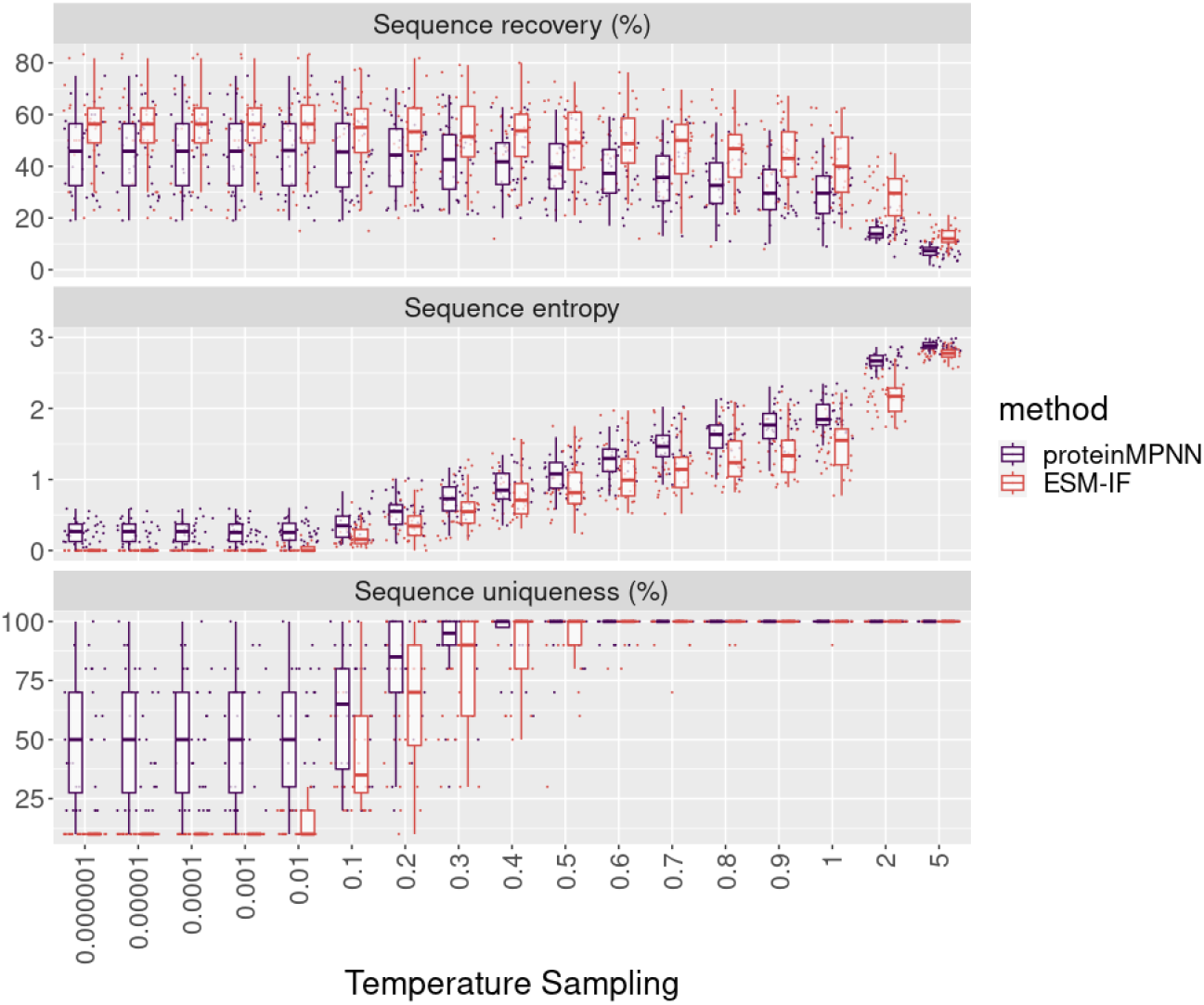
Effects of temperature sampling on sequence recovery, entropy and uniqueness in the design of interface CDR3s residues with ProteinMPNN and ESM-IF. The upper panel presents the sequence recovery relative to the temperature sampling. Each point corresponds to the average sequence recovery of each MHC-I test case. The middle panel presents the entropy of the designed sequences for each MHC-I test case in function of the temperature sampling. Higher entropy indicates higher diversity in the generated sequences. The entropy was calculated using the R Bio3d package as an average of positional entropies. The bottom panel shows the uniqueness of generated sequences in function of temperature. Maximum uniqueness (100%) indicates that all generated sequences for a given test case are different. The tested temperatures ranged from 0.000001 to 5.

**Figure S3:**
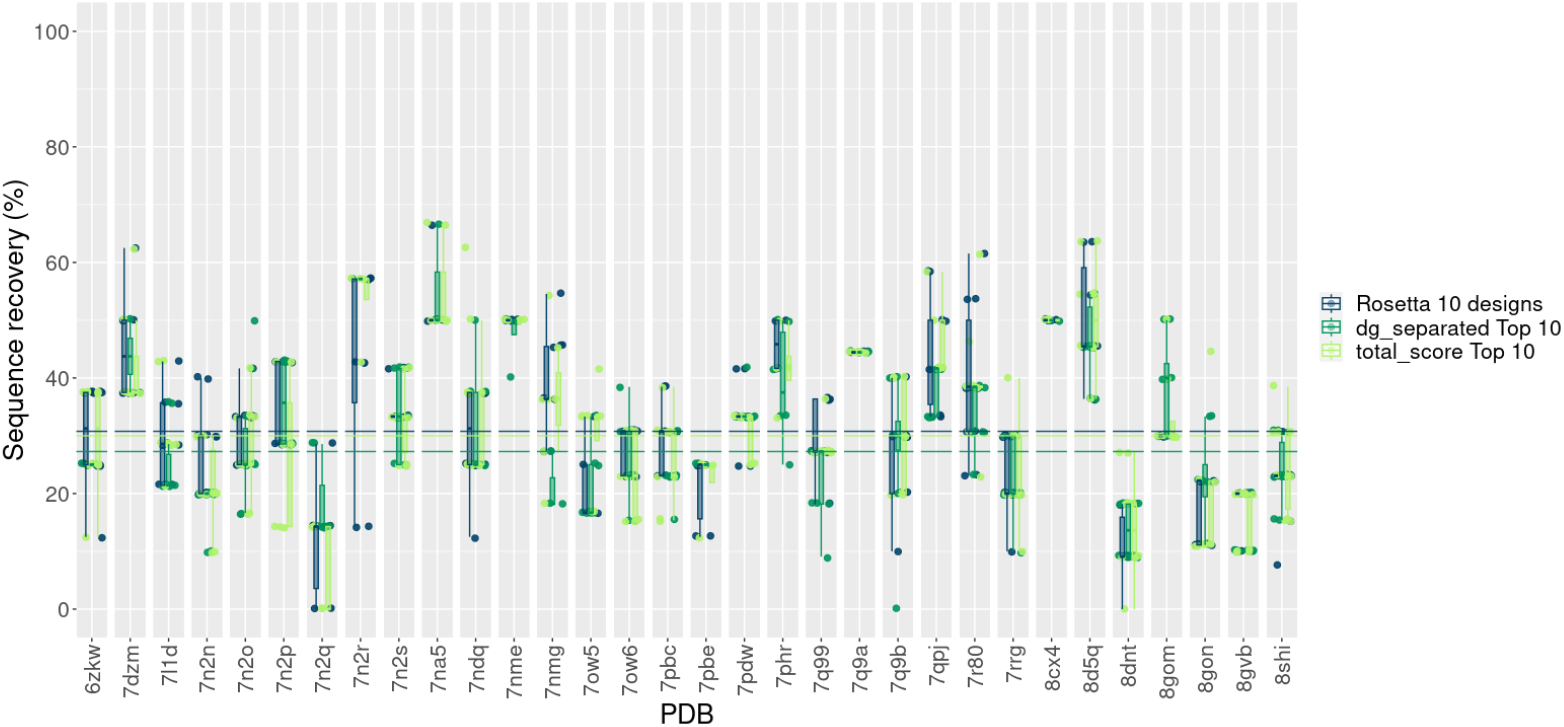
Sequence recovery analysis of interface CDR3 amino acids in designing with different flavors of Rosetta InterfaceDesign2019 protocol. Dark green box plots represent the sequence recovery for each target using the default Rosetta InterfaceDesign2019 protocol and generating 10 designs per target. The medium green box plots represent the sequence recovery of top 10 designs scored by Rosetta dg_separated term from a total of 1000 generated designs. The light box plots represent the sequence recovery of top 10 design scores by Rosetta total_score term from a total of 1000 generated designs. Each point corresponds to a design sequence. Redundant designs were removed from the analysis. Lines indicate the median computed over all designs: 30.8%, 27.3%, and 30.0% for 10 designs with default Rosetta protocol, top 10 best dg_separated score designs and top 10 best total_score scored designs, respectively.

**Figure S4:**
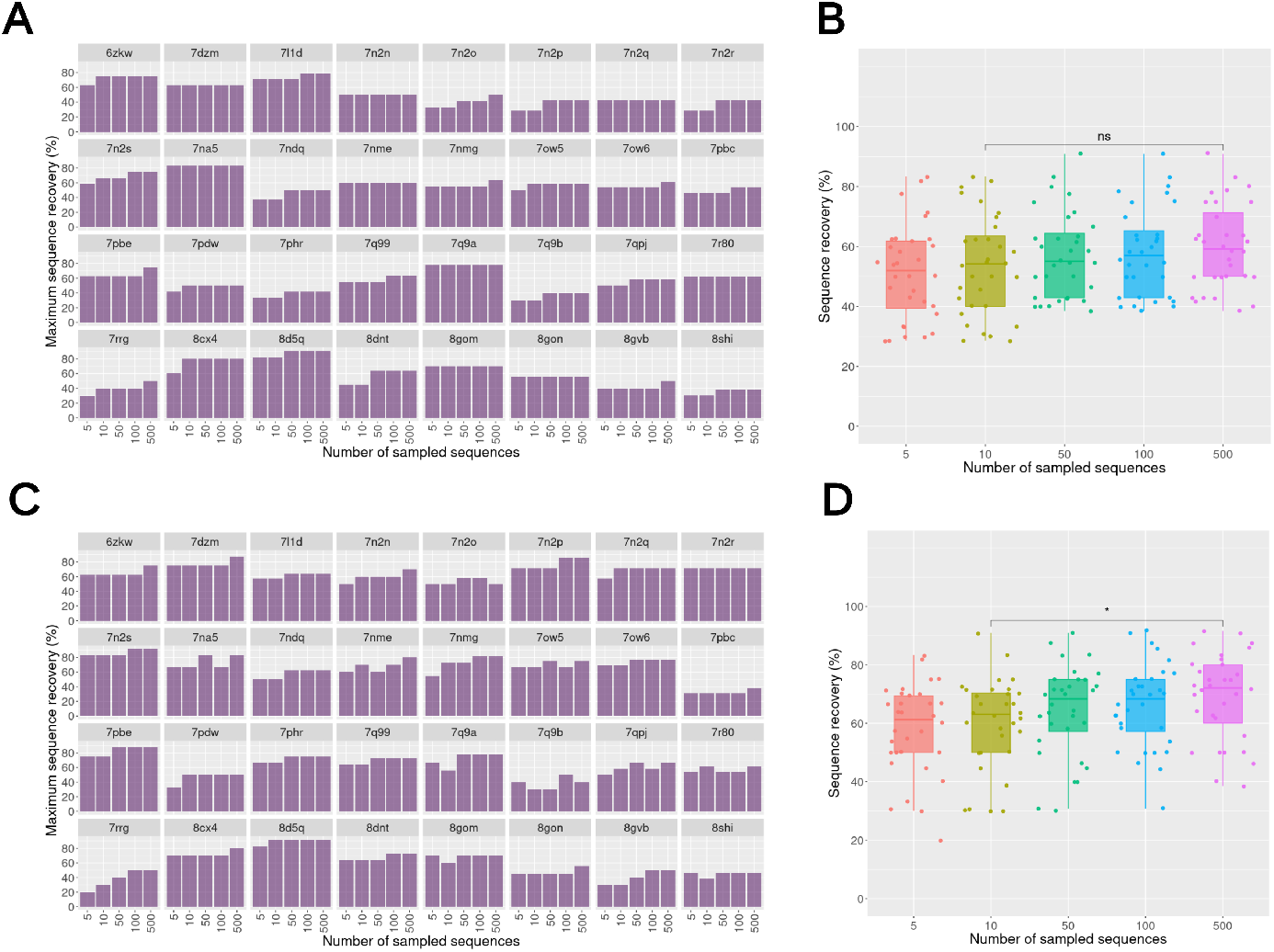
ProteinMPNN (A) and ESM-IF (B) TCR design considering different design scenarios. The box plots present the sequence recovery for each design strategy: design of only CDR3 (α and β chains) at the interface with the pMHC (in purple), design of all CDR3 (α and β chains) positions (in green) and design of CDR1, CDR2, and CDR3 positions (α and β chains) at the interface with the pMHC. Each point corresponds to the sequence recovery of a designed sequence. Redundant sequences within each test case were excluded from the analysis. In (A), lines indicate the median computed over all designs: 44.4%, 53.1%, and 39.0% for CDR3s interface positions, all CDR3 positions and all CDRs interface positions, respectively. In (B), lines indicate the median computed over all designs: 50%, 53.8%, and 45.3% for CDR3s interface positions, all CDR3 positions and all CDR interface positions, respectively.

**Figure S5:**
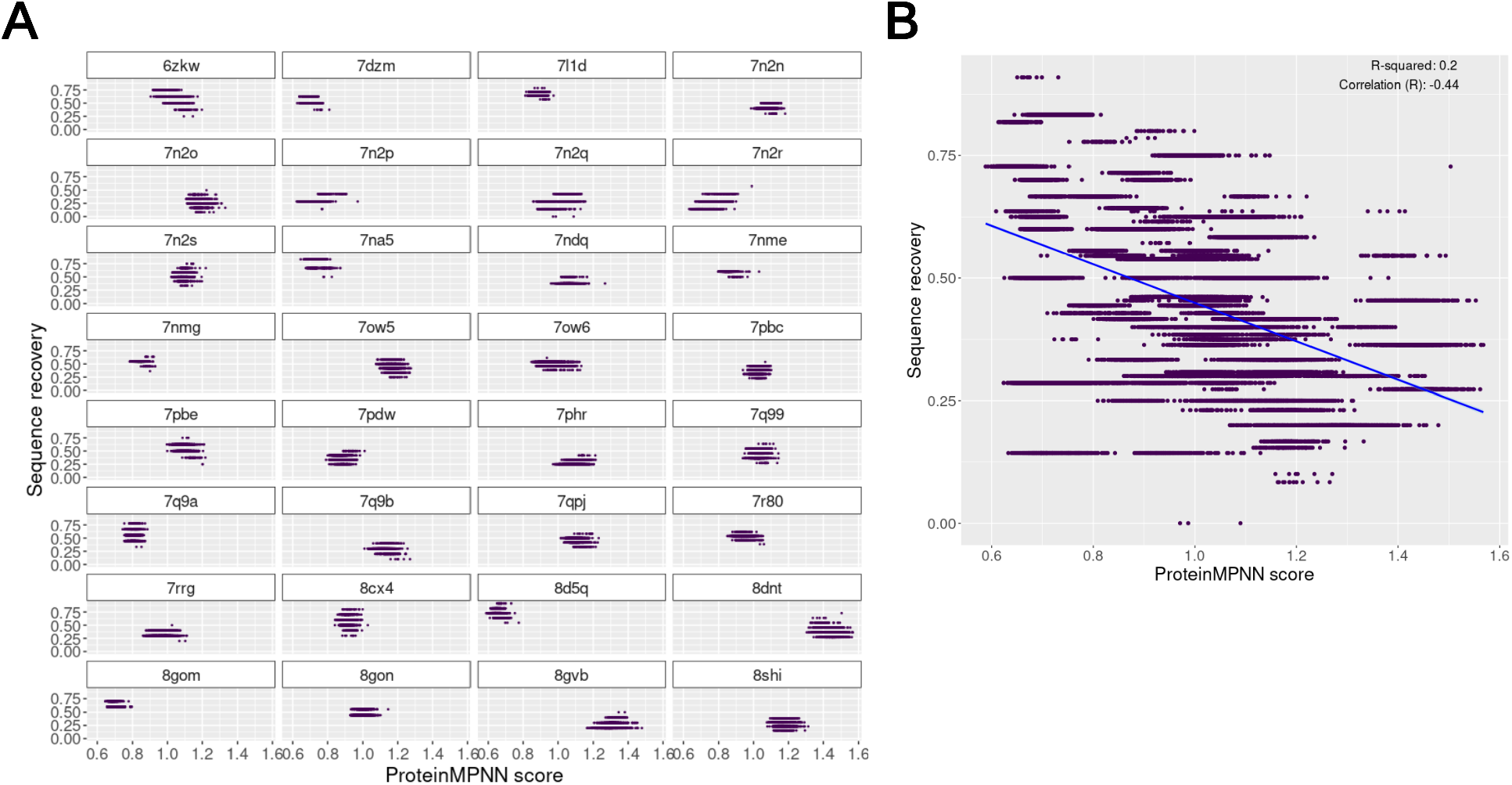
Correlation between ProteinMPNN sequence recovery and score. **(A)** Scatter plot of sequence recovery and ProteinMPNN scores from 1000 ProteinMPNN designs for each test case. Redundant sequences were not removed in this analysis since the same sequence can have different ProteinMPNN scores. **(B)** Scatter plot considering all designs together with a linear trend line (blue line) presenting the correlation between the sequence recovery and ProteinMPNN score. Correlation coefficients are indicated in the plot. The correlation coefficient was determined using Spearman.

**Figure S6:**
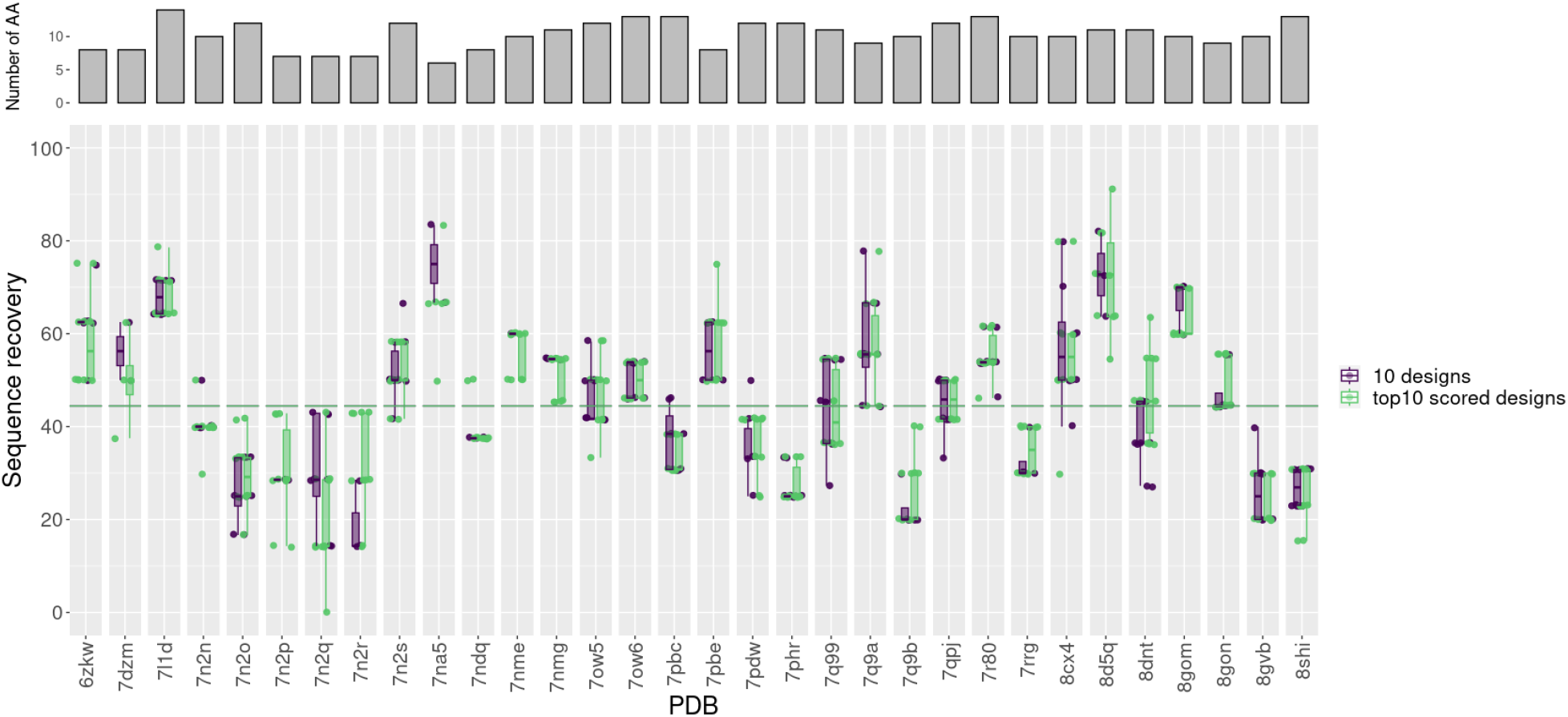
Comparison of the sequence recovery from 10 ProteinMPNN designs or a top 10 of designs ranked by ProteinMPNN score from a total of 1000 generated designs. The box plots present the sequence recovery for the 10 ProteinMPNN designs (in purple) and for the top 10 scored designs (in green). Each point corresponds to the sequence recovery of a designed sequence. Redundant sequences within each test case were excluded from the analysis. Lines indicate the median computed over all designs: 44.4% for both cases.

**Figure S7:**
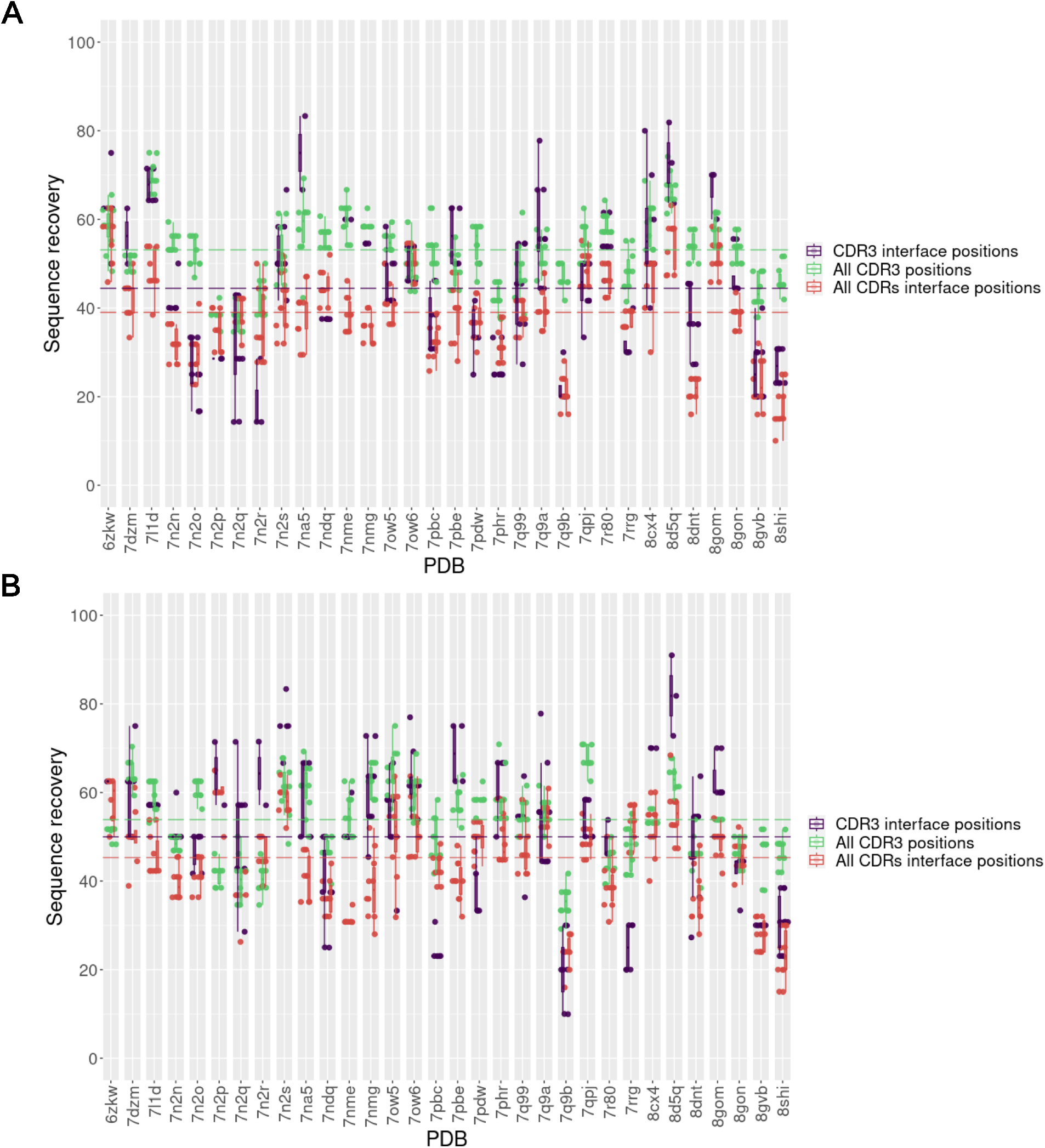
ProteinMPNN (A) and ESM-IF (B) TCR design considering different design scenarios. The box plots present the sequence recovery for each design strategy: design of only CDR3 (α and β chains) at the interface with the pMHC (in purple), design of all CDR3 (α and β chains) positions (in green), and design of CDR1, CDR2, and CDR3 positions (α and β chains) at the interface with the pMHC. Each point corresponds to the sequence recovery of a designed sequence. Redundant sequences within each test case were excluded from the analysis. In **(A)**, lines indicate the median computed over all designs: 44.4%, 53.1%, and 39.0% for CDR3s interface positions, all CDR3 positions, and all CDRs interface positions, respectively. In **(B)**, lines indicate the median computed over all designs: 50.0%, 53.8%, and 45.3% for CDR3s interface positions, all CDR3 positions and all CDR interface positions, respectively.

**Figure S8:**
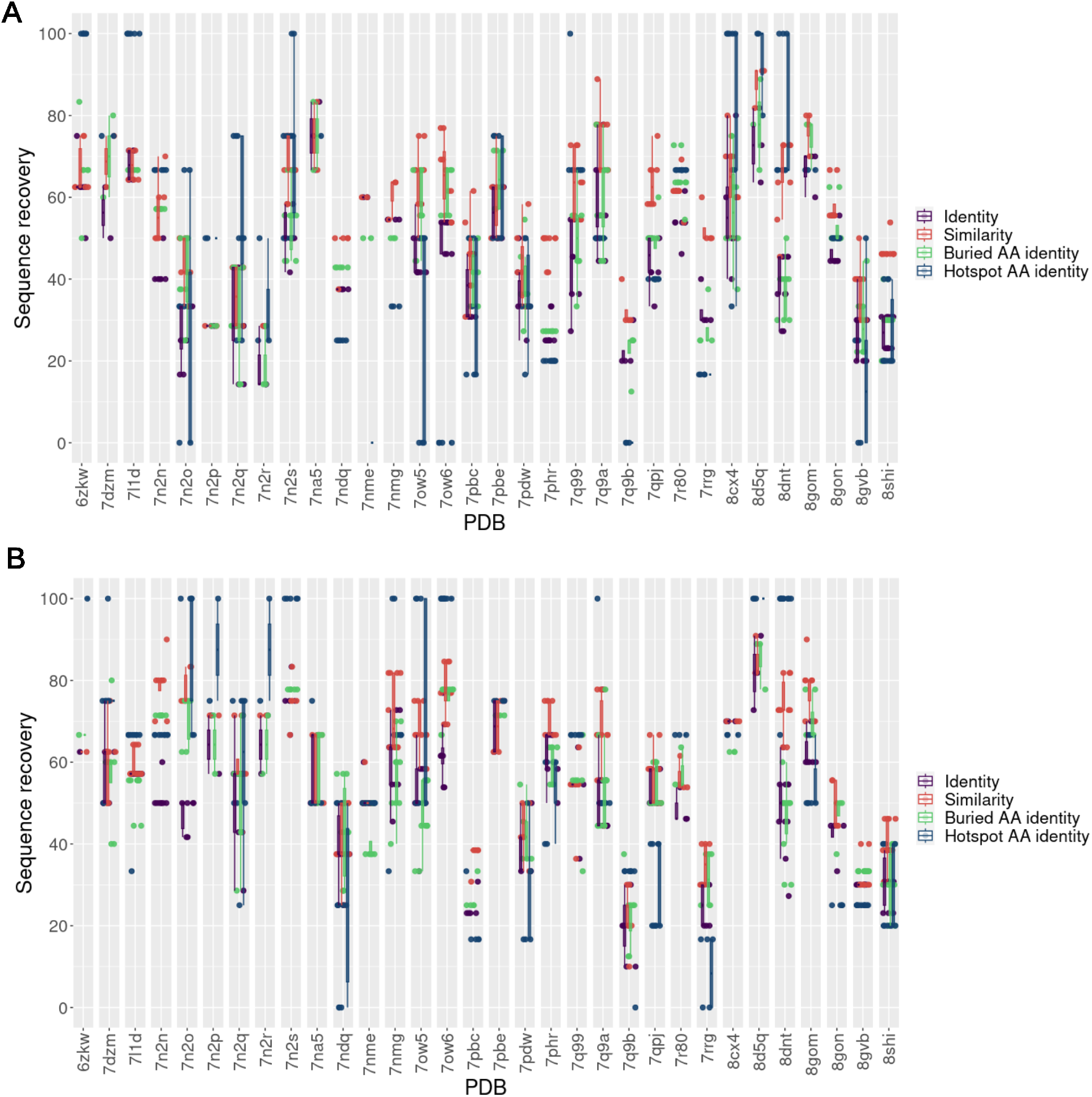
**Sequence recovery analysis of interface CDR3 amino acids in terms of identity (same amino acids in design and native sequence), similarity (amino acids in design within the same physicochemical group as the one in native sequence), identity at only interface buried positions and identity at hotspot positions for ProteinMPNN (A) or ESM-IF (B)**. The sequence recovery (in %) is presented by box plots for each case designed by ProteinMPNN. Each point corresponds to a design sequence.

**Figure S9:**
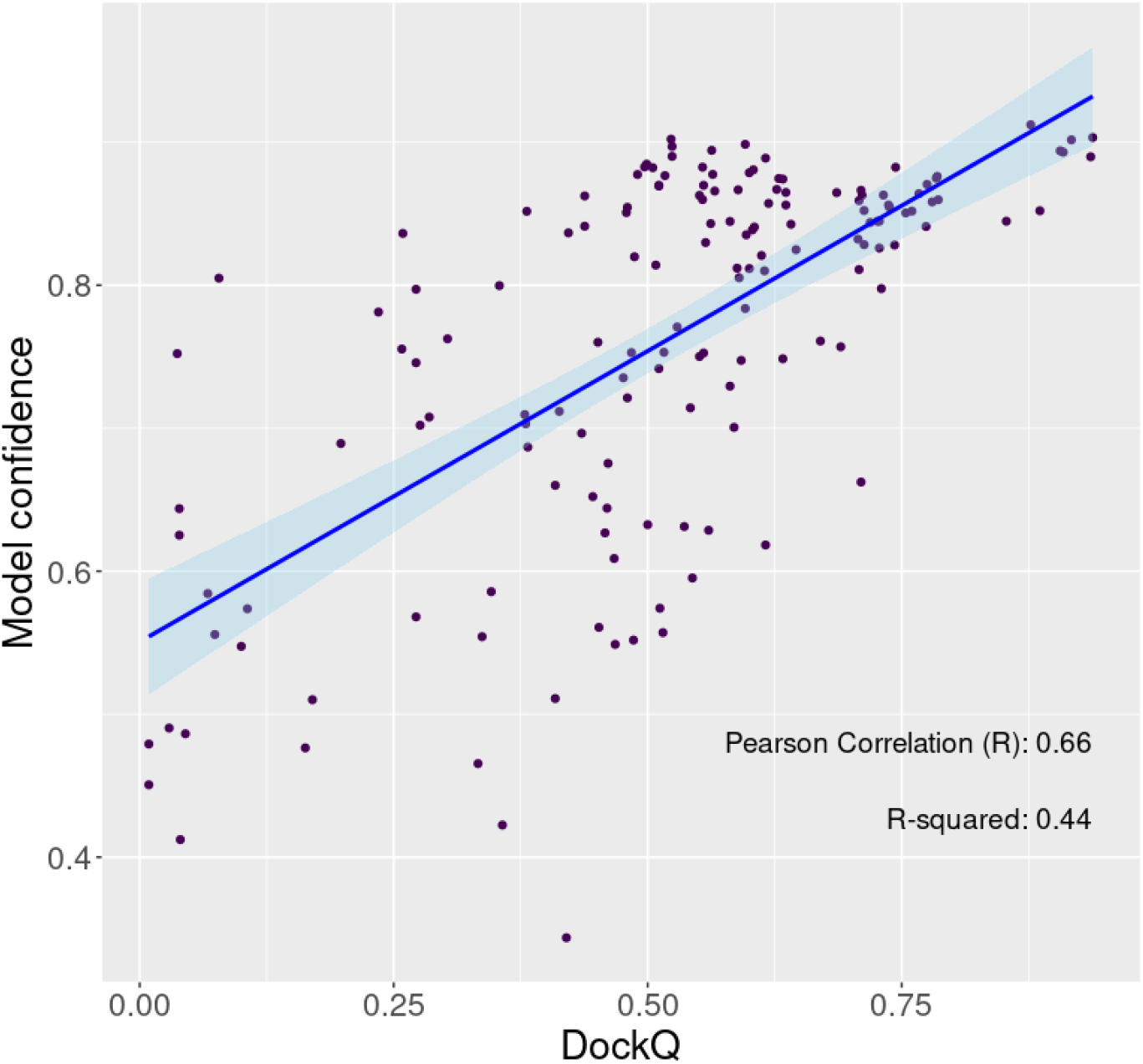
Correlation between model confidence and DockQ obtained for TCRModel2 models (5 models per each test case) of remodeled native sequences.

**Figure S10:**
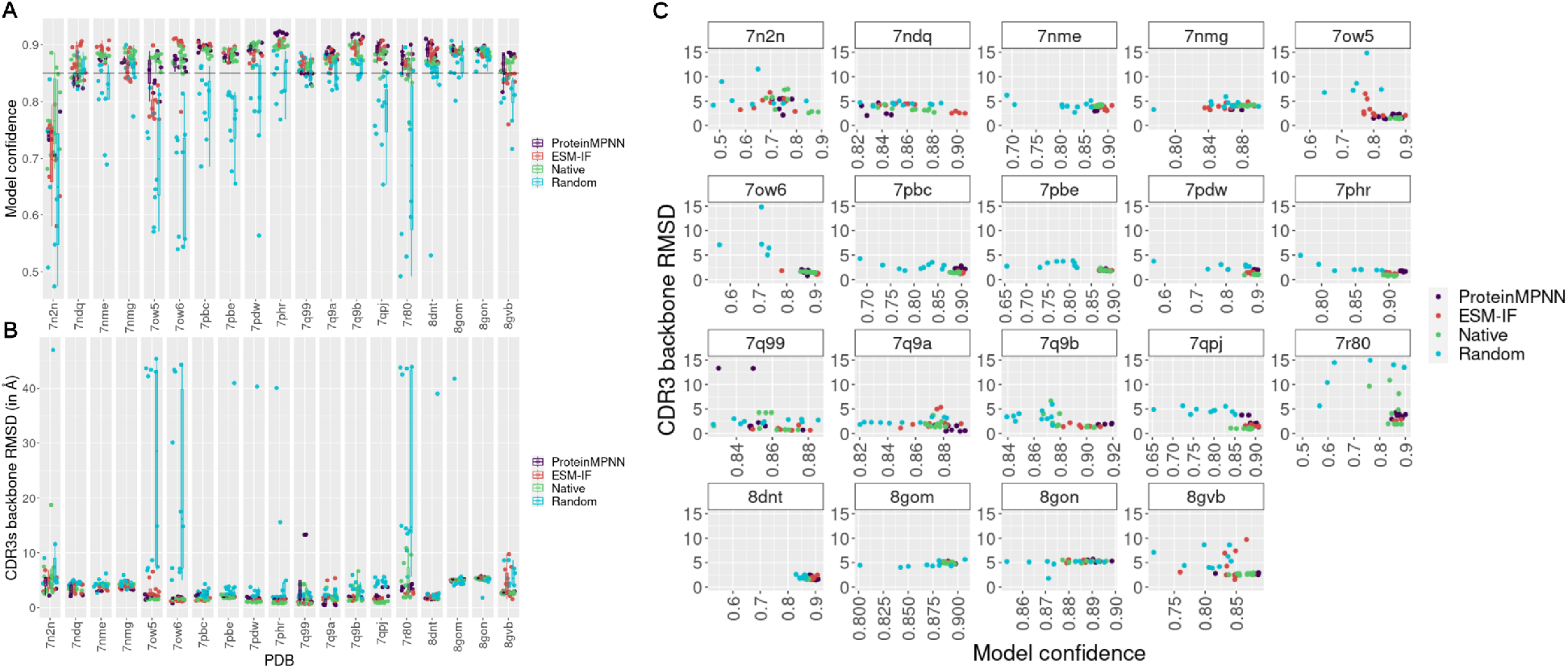
Modeling of ProteinMPNN and ESM-IF TCR designs, native and random dissimilar sequences with TCRmodel2. **(A)** Model confidence of ProteinMPNN (in purple), ESM-IF (in red) TCR designs, remodeled native structures (in green) and random dimissilar sequences (in cyan) for each test case. **(B)** RMSD of CDR3 backbone atoms (both alpha and beta TCR chains) of designs and random in comparison to the corresponding native crystal structure that originated the designs. RMSDs were determined after structure superposition by the MHC. **(C)** Scatter plot of the CDR3 backbone RMSD with the model confidence for ProteinMPNN and ESM-IF designs, random dissimilar sequences and native sequences. Since random sequences generate models with high deviation, for clarity only RMSD below 15 Å are presented.

**Figure S11:**
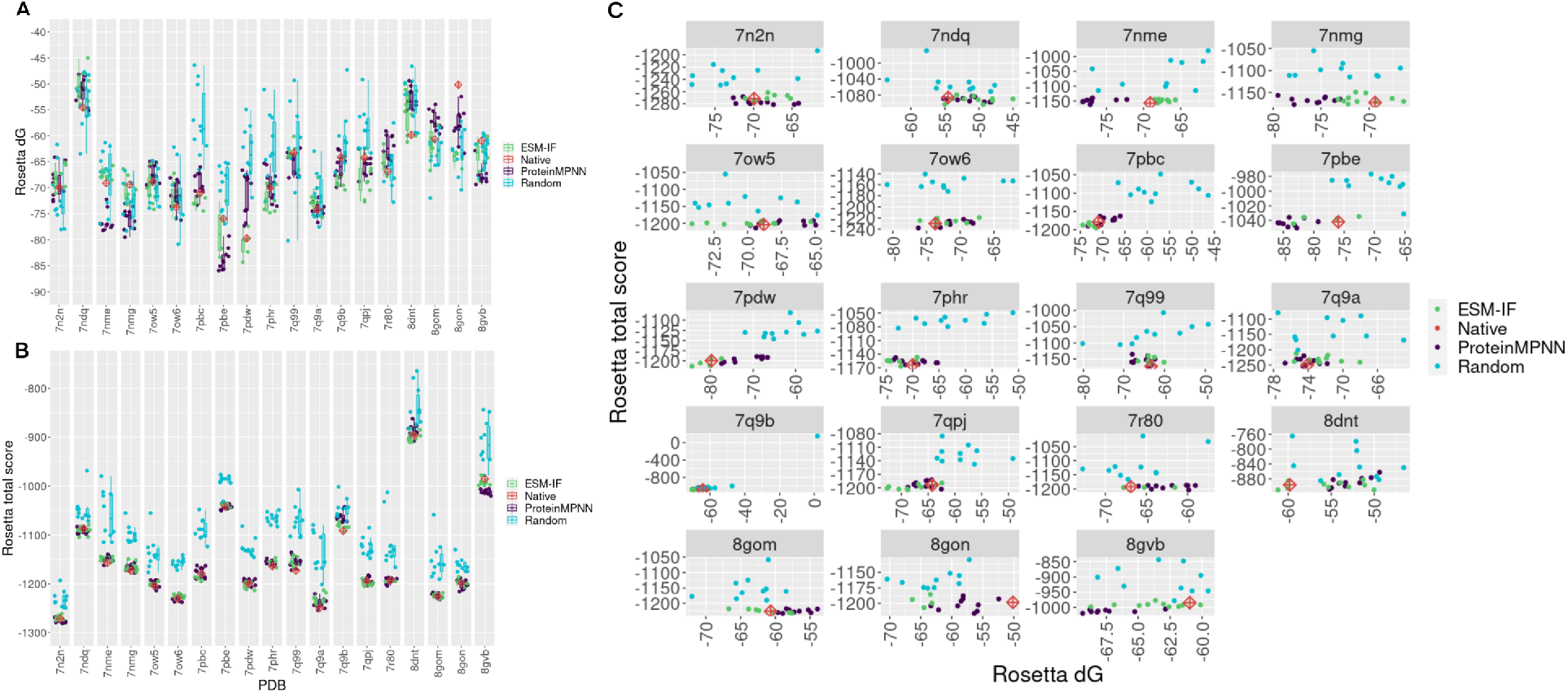
Scoring of ProteinMPNN and ESM-IF designs, native structure and dissimilar random generated structures with Rosetta energy functions. **(A)** Rosetta *dG_separated* term obtained from Rosetta InterfaceAnalyzer. Box plots represent *dG_separated* of the ESM-IF (in green) and ProteinMPNN (in purple) designs, dissimilar random generated TCRs (in cyan) and the red diamond corresponds to the *dG_separated* of the native structure. **(B)** Same as (A), but showing the Rosetta *total_score* term instead of *dG_separated*. **(C)** Scatter plot presenting the relation between the *dG_separated* and *total_score* for native, designs and random sequences for all test cases. The lower the *dG_separated* and *total_score* scores, the higher the affinity and stabilization, respectively.

**Figure S12:**
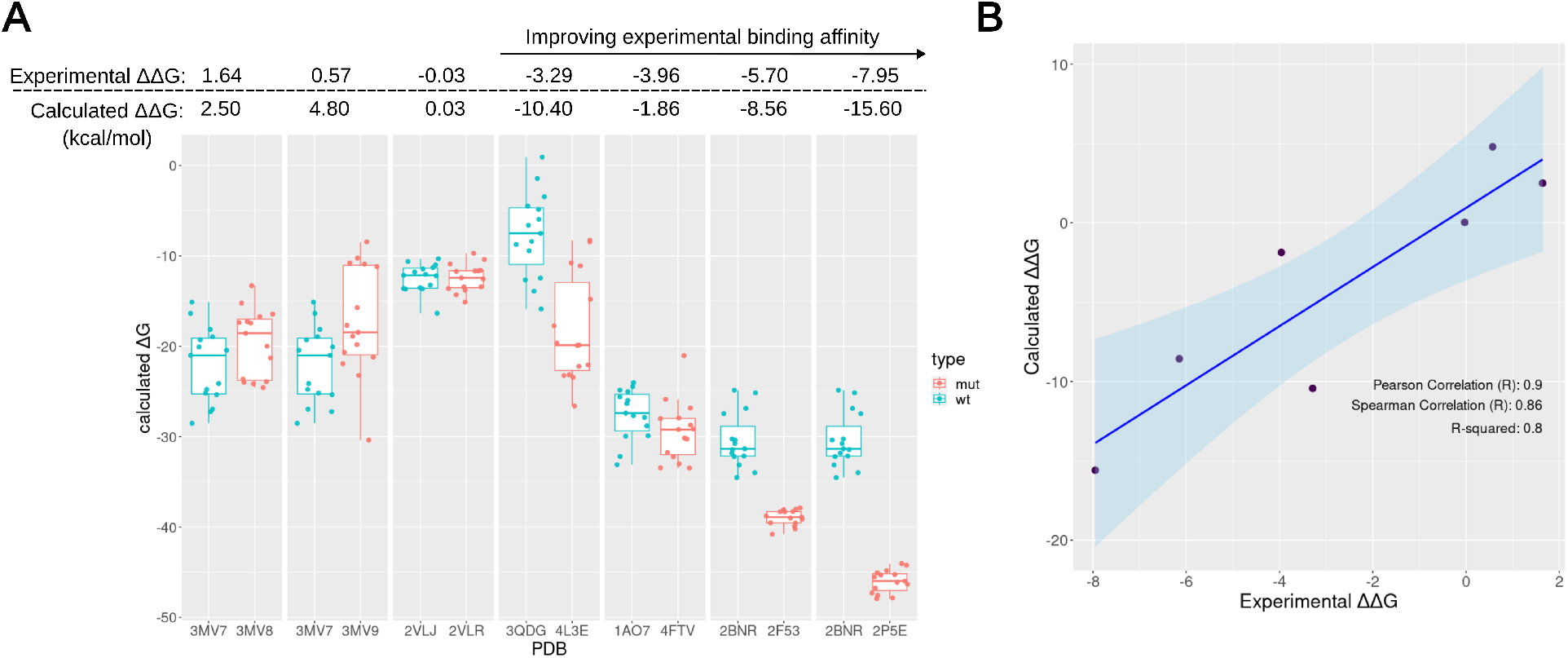
Binding affinity estimation of wild-type and mutant TCR complexes using molecular dynamics simulation and MM/PBSA calculations. **(A)** Box plots of calculated ΔG (in kcal/mol) for the wild-type TCR complex (wt, in blue) or mutant complex (mut, in red) from MM/PBSA. Each point corresponds to a replicate (15 in total) of a molecular dynamics simulation trajectory. The values of the experimental and calculated ΔΔG (ΔGmut - ΔGwt) (in kcal/mol) are shown above the box plot panel. **(B)** Scatter plot with a linear trend line and confidence interval of 0.95 (light blue region) presenting the correlation between the experimental and calculated ΔΔG. Correlation coefficients are indicated in the plot.

**Figure S13:**
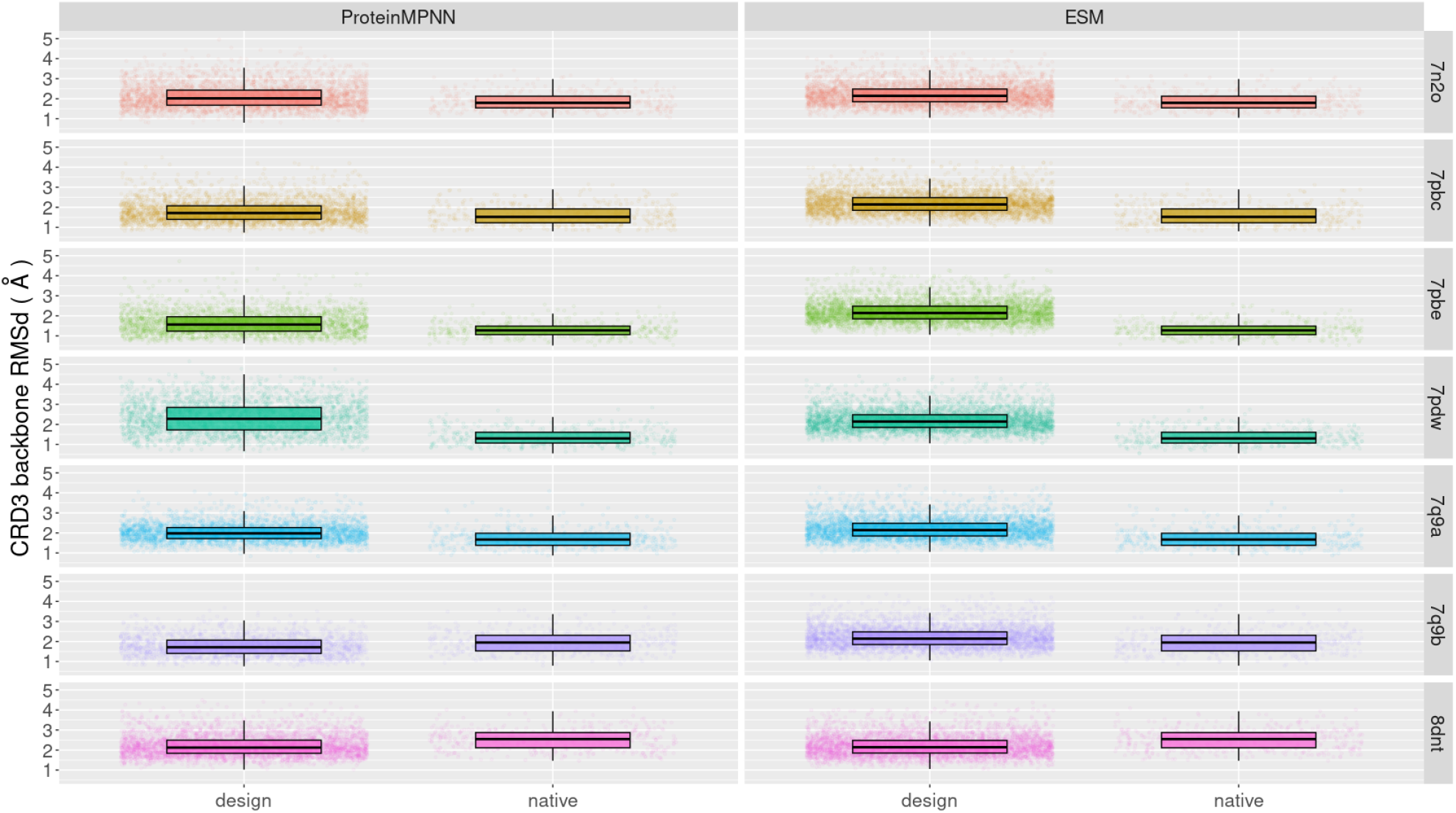
RMSD of CDR3 (α and β chains) backbone estimated from trajectories of molecular dynamics simulations of ProteinMPNN and ESM-IF designs, as well as the native trajectory. The RMSD values, depicted as box plots, were calculated subsequent to superposing the trajectory frames by the MHC of the corresponding reference crystal structure. For each test case, all replicas of all designs were combined to form a single RMSD distribution.

**Figure S14:**
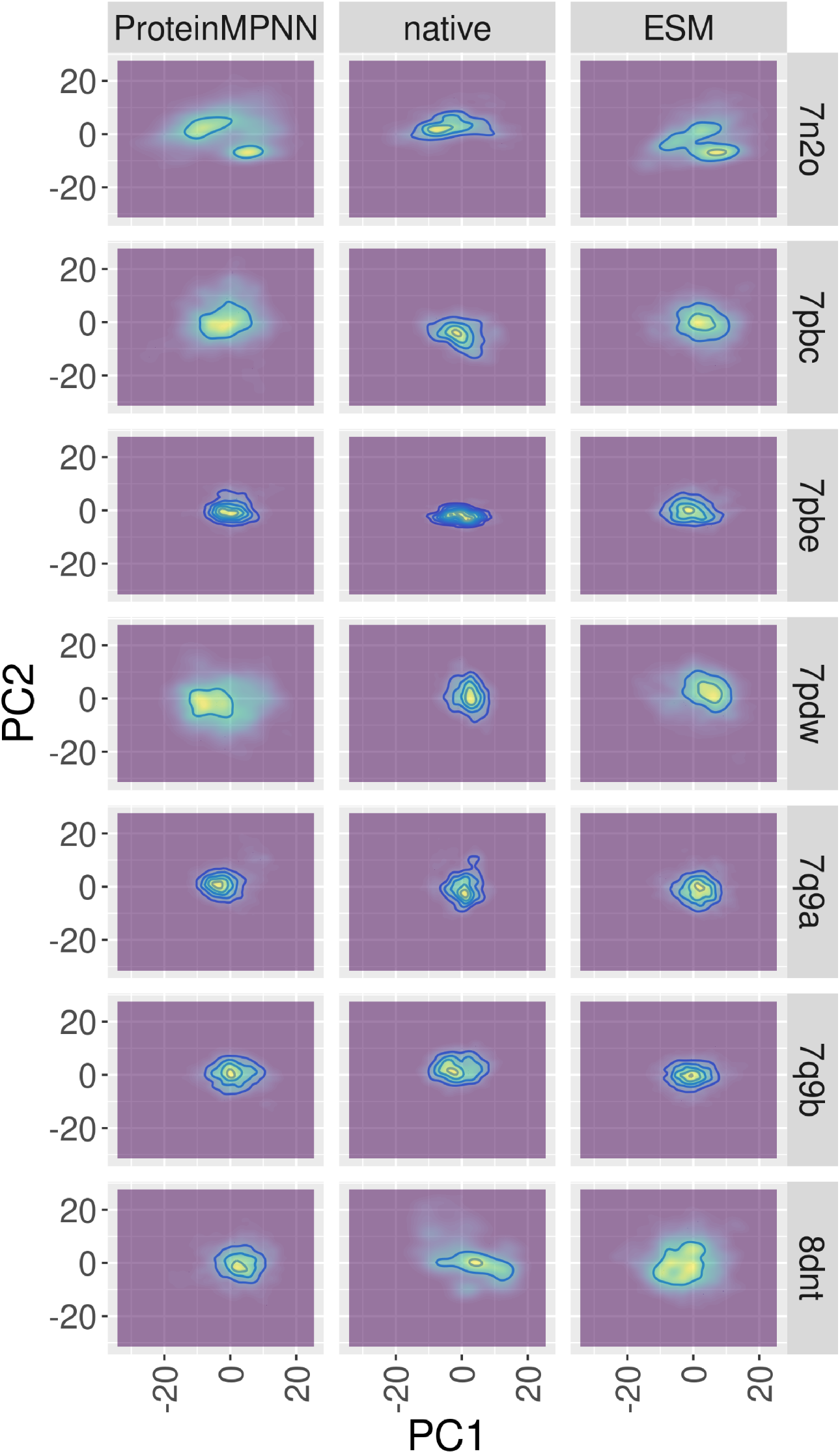
Principal Component Analysis (PCA) of molecular dynamics simulations of ProteinMPNN and ESM-IF designed TCRs and native TCRs. For the PCA analysis, only the CDR3 coordinates were considered (see methods). The first two main components are represented as 2D density contours (bins of 50) colored by R viridis scale that ranges from dark purple to yellow, being yellow the regions of higher density. Plots were built using R ggplot package.

**Figure S15:**
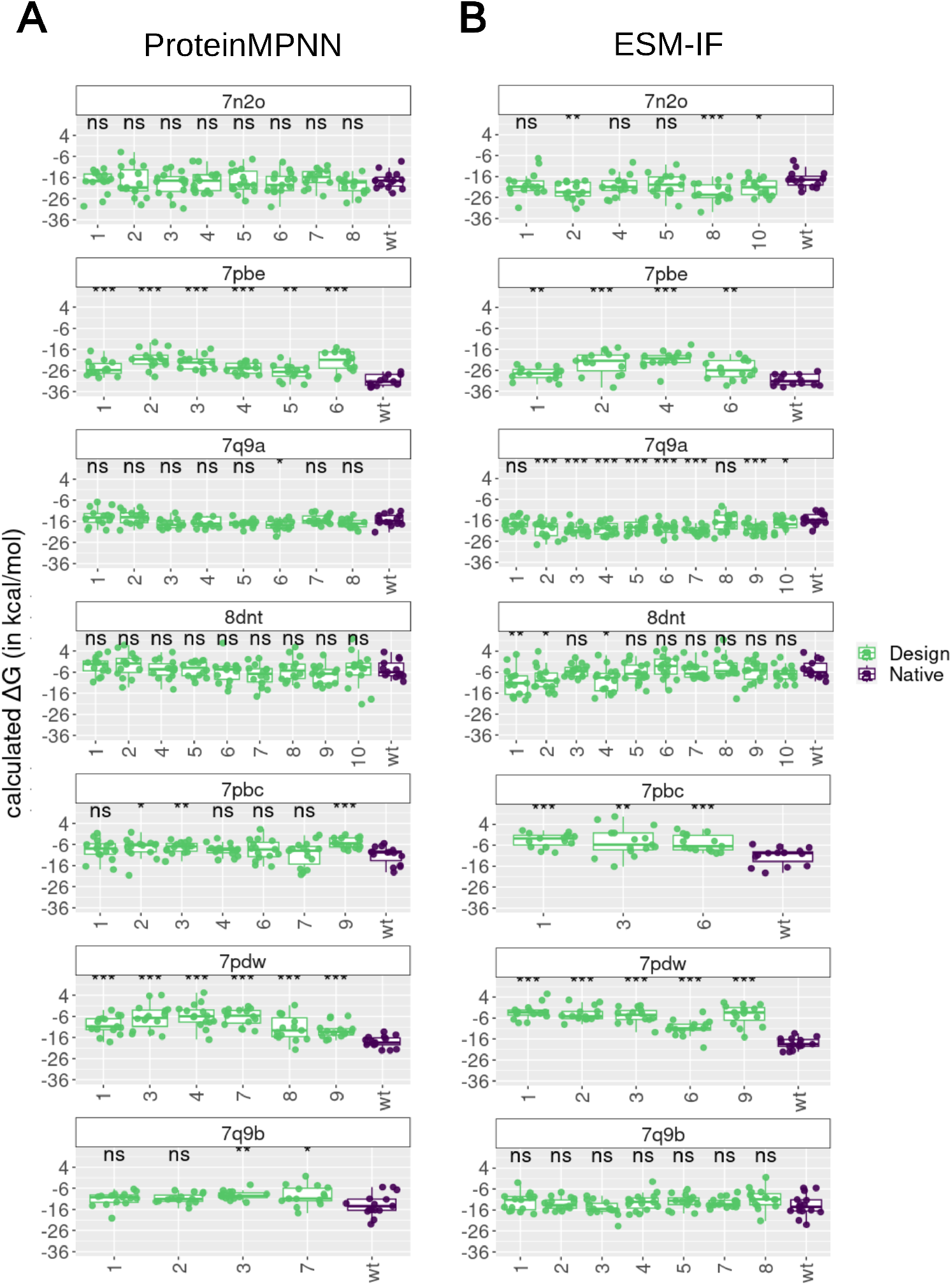
Binding affinity estimation of ProteinMPNN (A) and ESM-IF (B) TCR designs by MM/PBSA calculations. Each plot corresponds to a TCR:pMHC test case. The box plots present the ΔG (in kcal/mol) calculated for each of the 15 replicas of the TCR designs (in green) and the wild-type (wt) TCR (in purple). The TCR designs are presented by IDs with a maximum of 10 designs. The lower number of designs are a consequence of redundant generated designs that were removed for the calculations. The statistical difference between each design and the corresponding wild-type was determined by Mann-Whitney test and the significance is indicated above each box plot (***, ** and * correspond to a p-value below 0.001, 0.01, and 0.05, respectively, while ‘ns’ means no significance).

**Figure S16:**
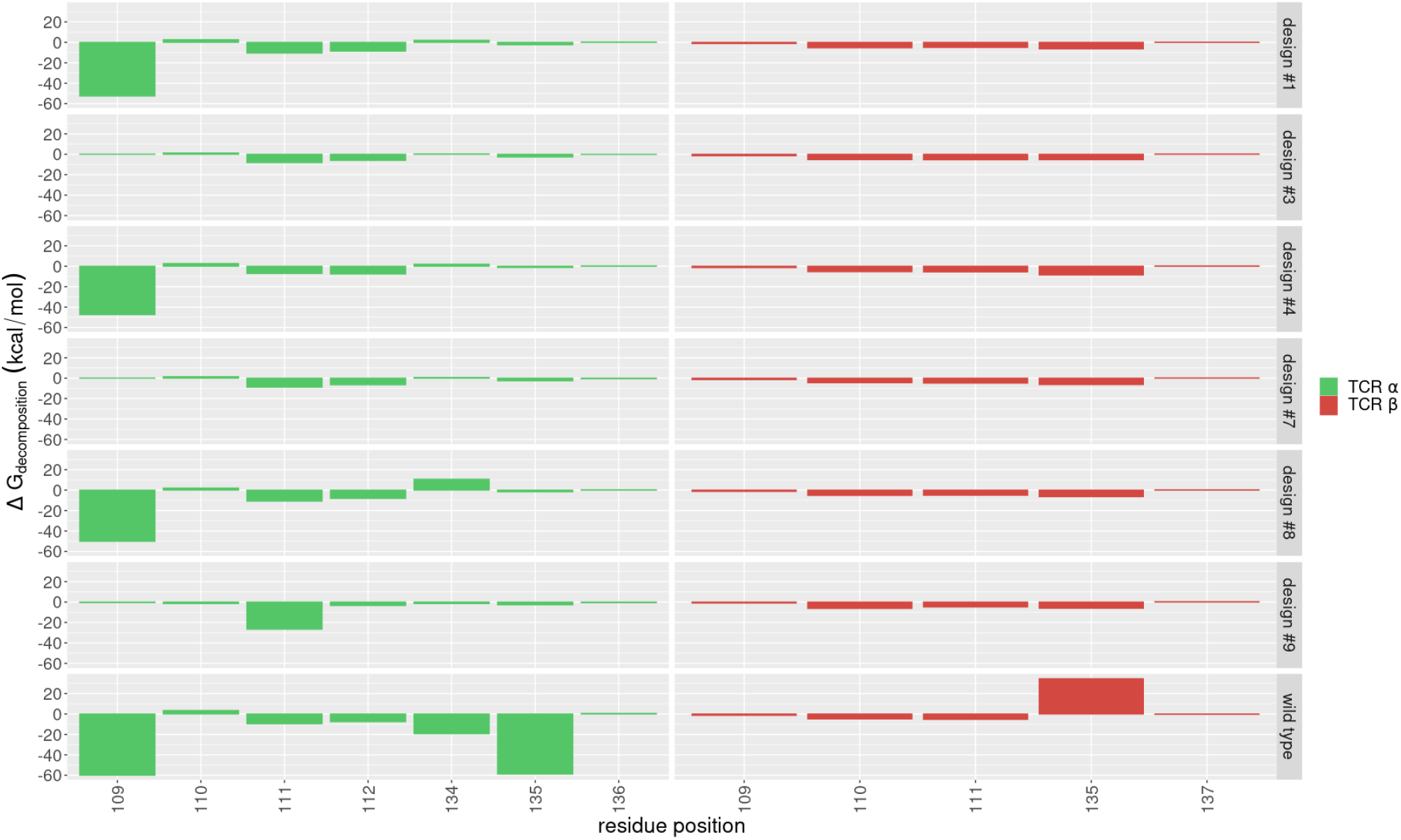
MM/PBSA decomposition (in terms of ΔG, in kcal/mol) of the TCR residues included in the design. Residues from TCRα (green) and TCRβ (red) are shown. Low ΔG values indicate the residue contributed positively to the binding energy and high ΔG values indicate the residue contributed negatively to the binding energy.

## B Supplementary Tables

**Table S1:**
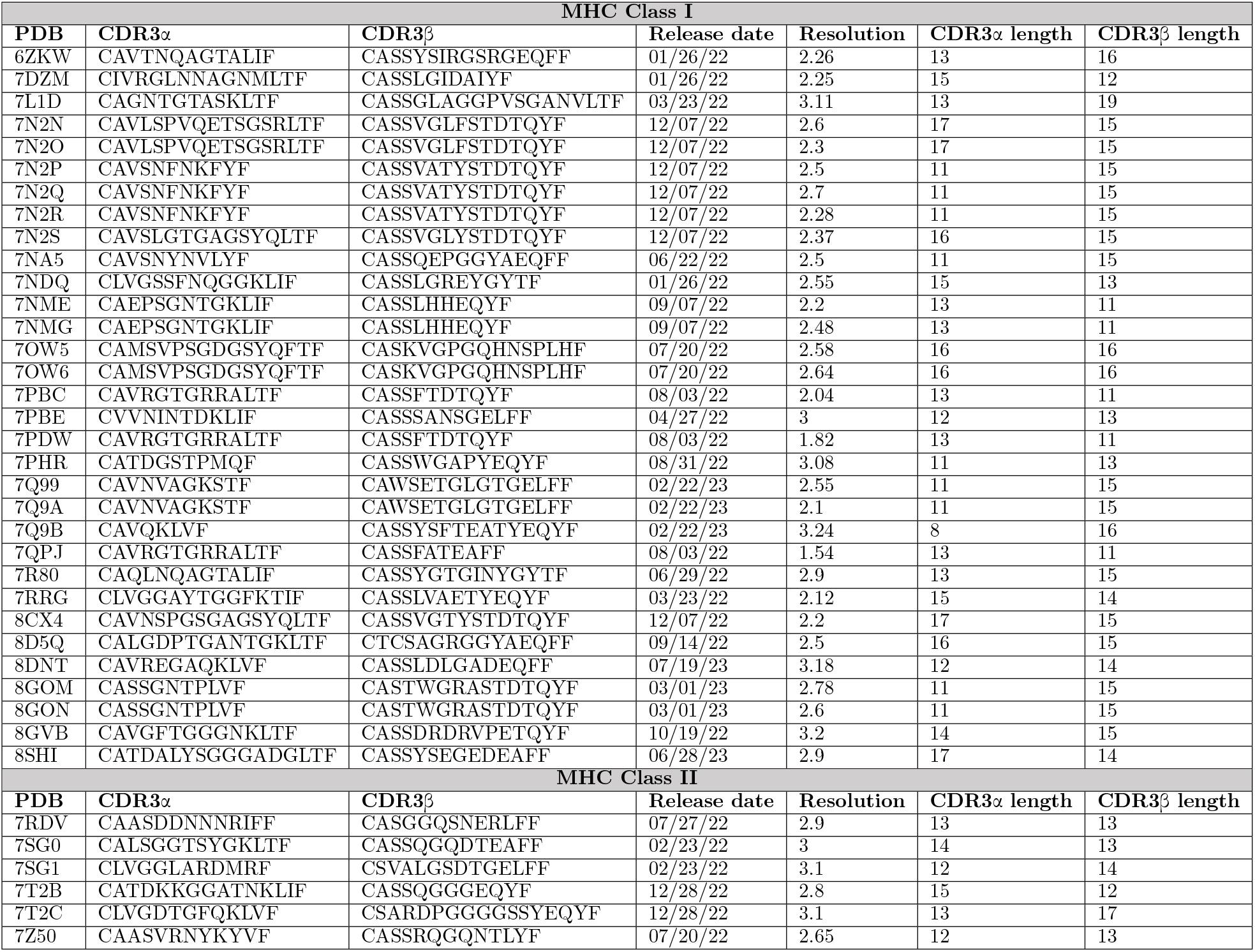
List of PDB IDs of MHC-I and MHC-II TCR:pMHC complexes used for comparing design methods.

**Table S2:**
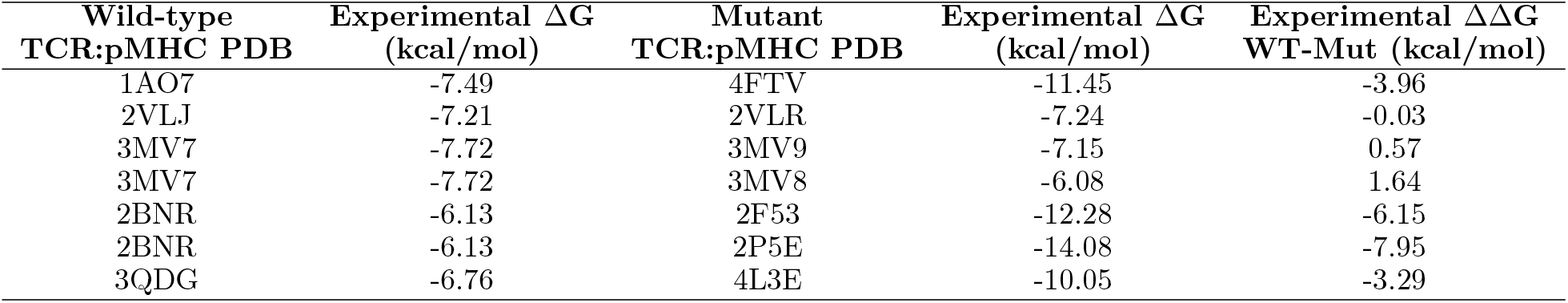
List of the wild-type and the corresponding mutant PDB structures of TCR:pMHC complexes that compose the benchmark for binding affinity calculations with MM/PBSA. The table includes the experimental ΔG (in kcal/mol) for each complex and the corresponding ΔΔG (ΔG_*mut*_ - ΔG_*wt*_). All experimental data was collected from ATLAS database (https://atlas.wenglab.org/).

## C Supplementary Information

### C.1 Rosetta resfile example

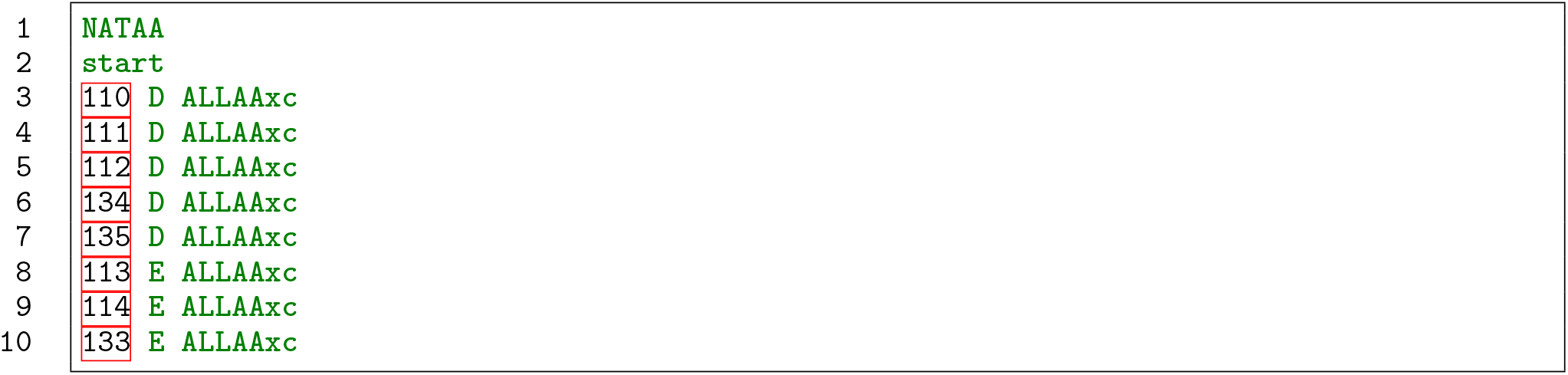

### C.2 Rosetta design protocol

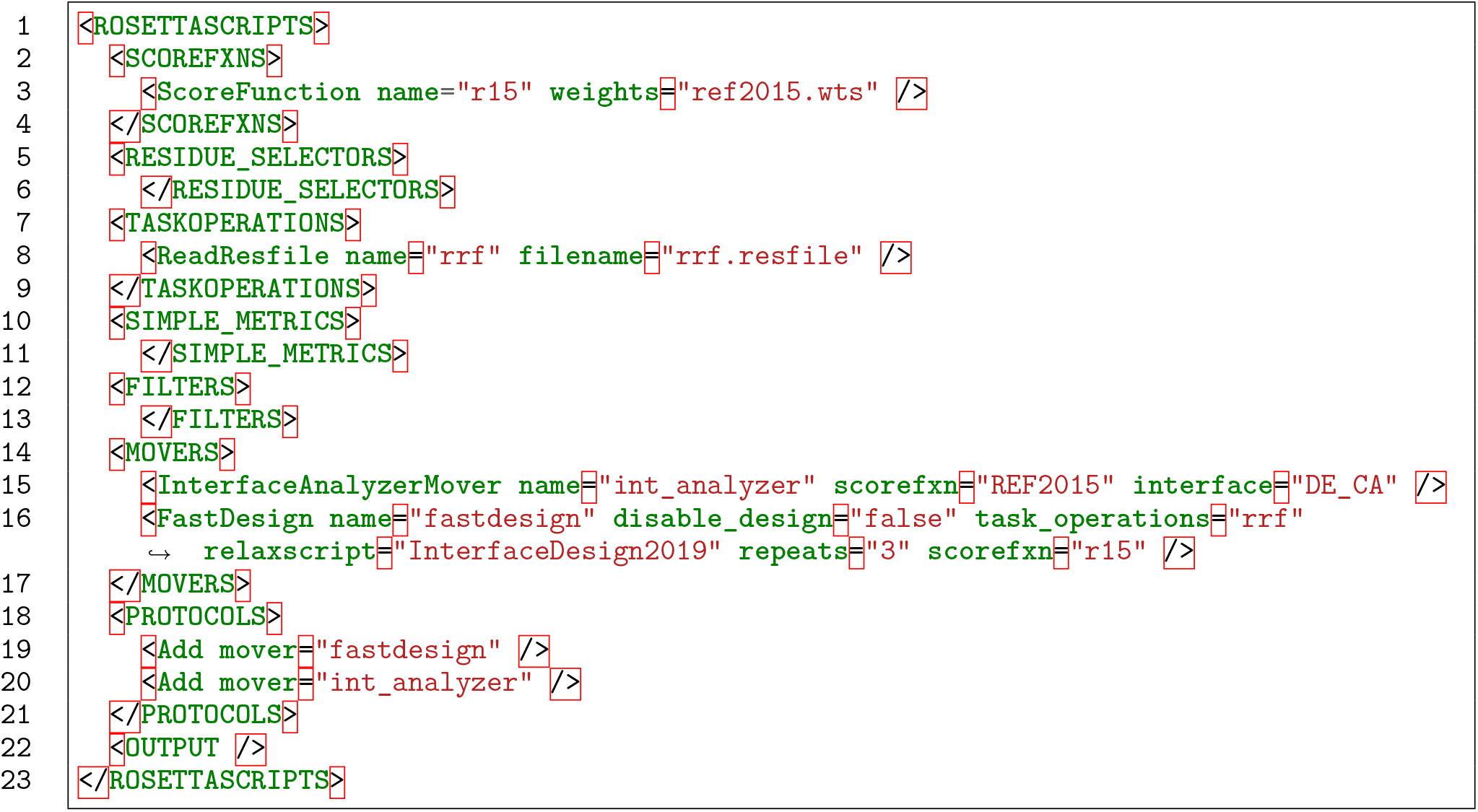

### C.3 Rosetta2.3 command line for alanine scanning

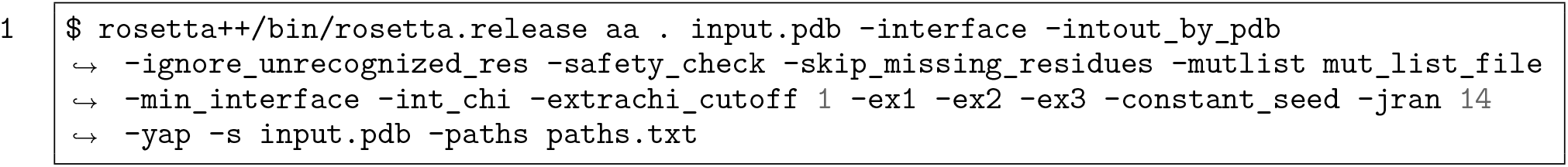

### C.4 Example of mutation list file for Rosetta2.3 alanine scanning protocol

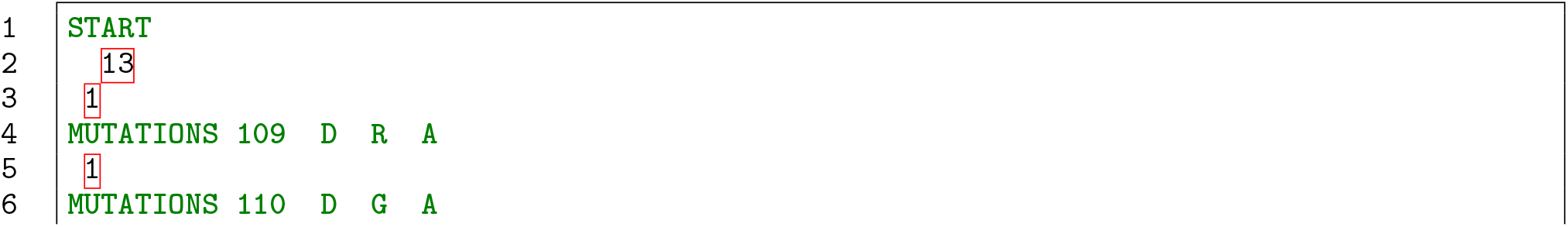

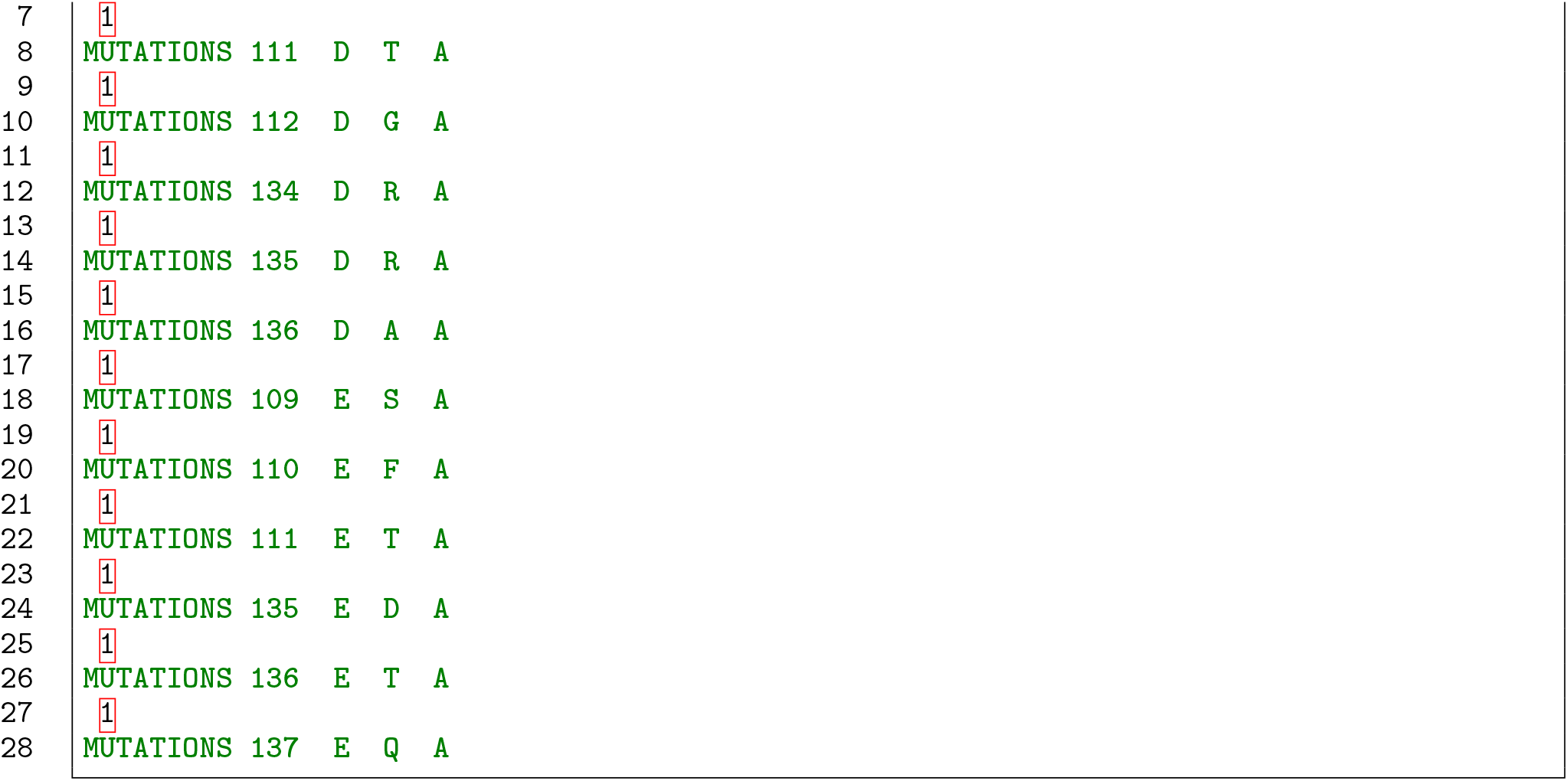

### C.5 Rosetta interface analyzer protocol

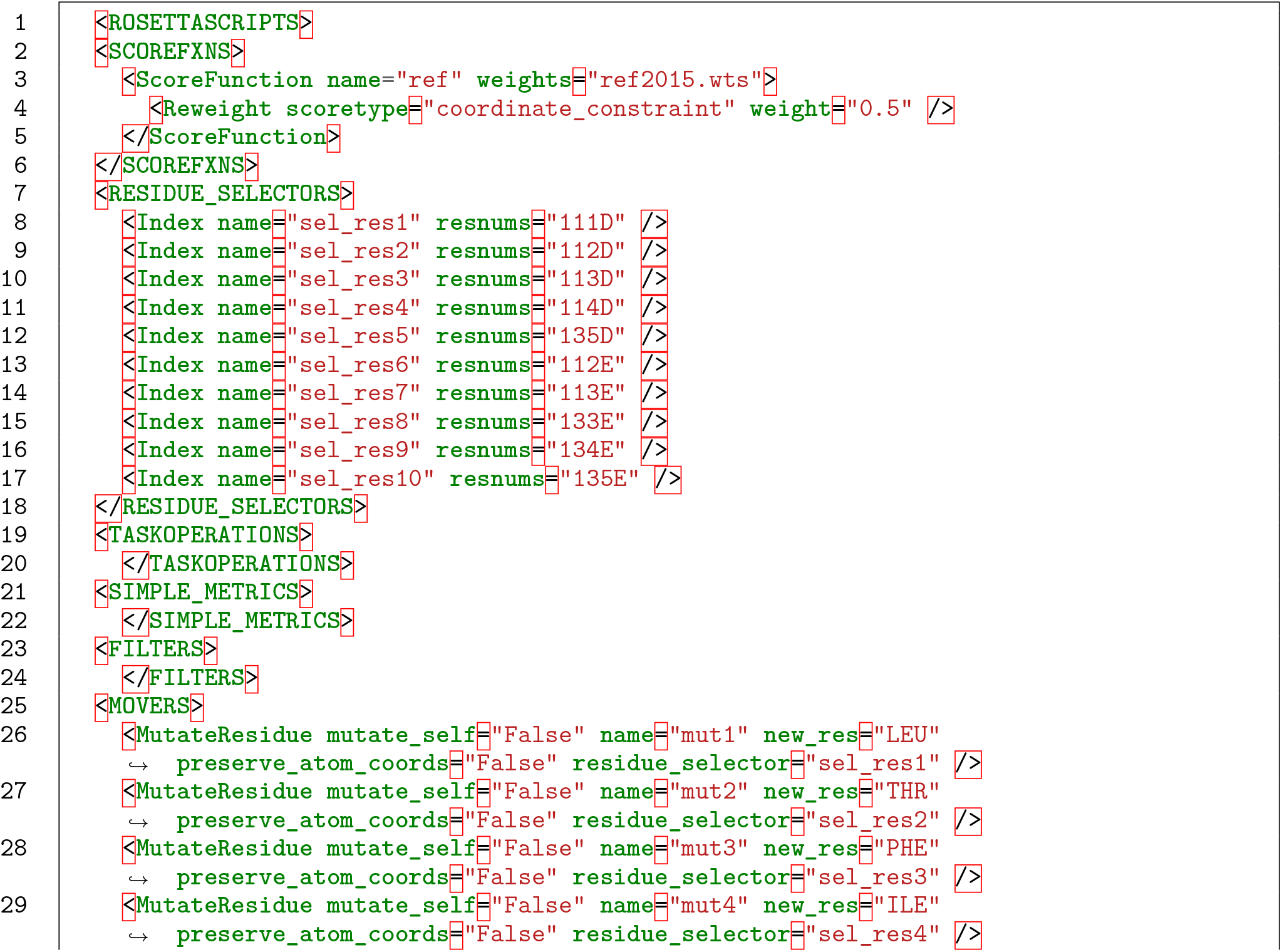

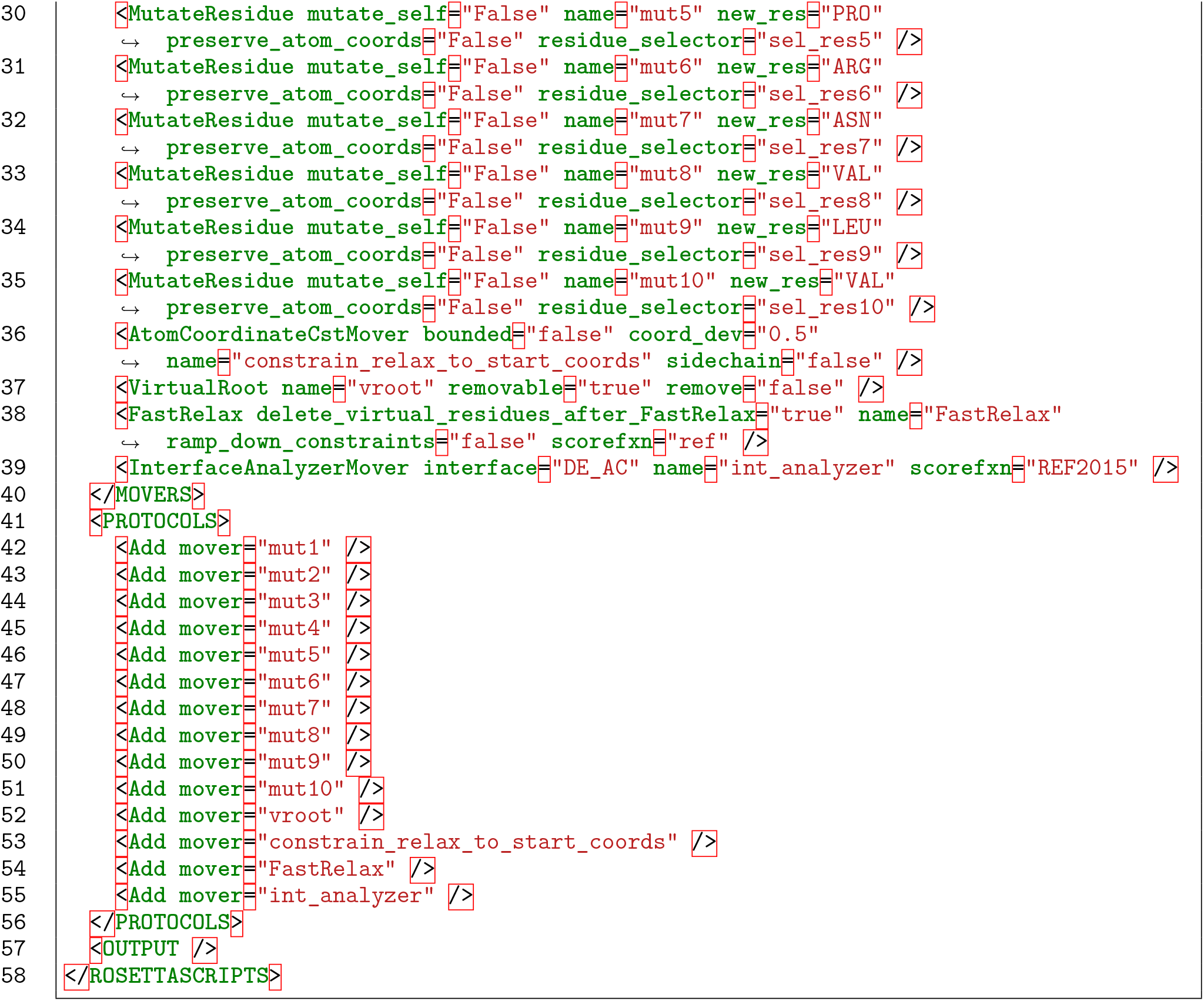

## Notes

### Competing Interest Statement

The authors have declared no competing interest.

